# β-bursts over frontal cortex track the surprise of unexpected events in auditory, visual, and tactile modalities

**DOI:** 10.1101/2022.07.13.499837

**Authors:** Joshua R. Tatz, Alec Mather, Jan R. Wessel

## Abstract

One of the fundamental ways in which the brain regulates and monitors behavior is by making predictions about the sensory environment and adjusting behavior when those expectations are violated. As such, surprise is one of the fundamental computations performed by the human brain. In recent years, it has been well-established that one key aspect by which behavior is adjusted during surprise is inhibitory control of the motor system. Moreover, since surprise automatically triggers inhibitory control without much proactive influence, it can provide unique insights into largely reactive control processes. Recent years have seen tremendous interest in burst-like β frequency events in the human (and non-human) local field potential – especially over (pre)frontal cortex) – as a potential signature of inhibitory control. To date, β-bursts have only been studied in paradigms involving a substantial amount of proactive control (such as the stop-signal task). Here, we used two cross-modal oddball tasks to investigate whether surprise processing is accompanied by increases in scalp-recorded β-bursts. Indeed, we found that unexpected events in all tested sensory domains (haptic, auditory, visual) were followed by low-latency increases in β-bursting over frontal cortex. Across experiments, β-burst rates were positively correlated with estimates of surprise derived from Shannon’s information theory, a type of surprise that represents the degree to which a given stimulus violates prior expectations. As such, the current work clearly implicates frontal β-bursts as a signature of surprise processing. We discuss these findings in the context of common frameworks of inhibitory and cognitive control after unexpected events.

## INTRODUCTION

Throughout the day, we constantly encounter external information translated by our various senses. While navigating this diverse sensory environment, our brain steadily generates predictions about this incoming stream of crossmodal information (Egner et al., 2010; Phillips et al., 2016). One of the key roles of frontal cortex is to detect deviations from such sensory predictions (Corbetta & Shulman, 2002; Kim, 2014). Indeed, identifying and modifying behavior according to unexpected perceptual events can have profound consequences on our lives. For instance, seeing a predator, hearing a roar, or feeling something crawl on one’s skin are all unexpected events that require rapid behavioral and cognitive adjustments. Subsequently, unexpected events provide a useful backdrop with which to explore the processes underlying adaptive behavior.

Indeed, unexpected perceptual events trigger multiple well-established cognitive and motor processes. Friston’s (2010) influential free-energy principle suggests that descending neural volleys convey predictions about the sensory environment, whereas ascending volleys convey violations of these expectancies, or prediction error. Some work has used simple mathematical models (Shannon, 1948; Baldi & Itti, 2010) to quantify surprise at the single-trial level, thereby showing that various neural signals may reflect such straightforward computations (e.g., Mars et al., 2008; O’Reilly et al., 2013; Nassar et al., 2019; Wessel & Huber, 2019). In addition to surprise processing, unexpected events are thought to capture attention in a bottom-up fashion and may interrupt goal-directed attentional representations, as in Corbetta and Shulman’s (2002) “circuit breaker” model of attention. Along these lines, target detection ability (e.g., Asplund et al., 2010), attentional representations indexed by steady-state visual evoked potential (e.g., Soh & Wessel, 2021), and content in active working memory may all be impaired by unexpected events (e.g., Wessel, 2018a; Hakim et al., 2021). Likewise, unexpected events prompt inhibition and disrupt ongoing action. Task-irrelevant unexpected events can induce reaction time slowing (e.g., Parmentier et al., 2008) as well as disrupt repetitive finger tapping (Horstmann, 2015) or the maintenance of isometric force contractions (Novembre et al., 2018). In addition, numerous studies using TMS to measure cortico-motor excitability have found a broad physiological suppression of the cortico-motor system after unexpected events, including at entirely task-irrelevant muscles (Wessel & Aron, 2013; Dutra et al., 2018; Iacullo et al., 2020; Tatz et al., 2021).

Further underscoring the link between surprise and inhibitory control, unexpected events and action cancellation are accompanied by common neurophysiological signatures. As the gold-standard method of testing action cancellation, the stop-signal task (SST) is the primary tool used to study motor inhibition (Verbruggen et al., 2019). In the SST, prepotent responses to Go signals must be inhibited upon interruption by an infrequent Stop signal. One common signature that is observed both after such stop-signals and after unexpected events outside of stop-signal paradigms is the above-mentioned suppression of the motor system (Badry et al., 2009; Cai et al., 2012; Wessel et al., 2013; Tatz et al., 2021). Another common signature is the fronto-central (FC) P3 event-related potential. The FC P3 ERP accompanies unexpected events and stop signals and is likely generated by the same underlying neural generator (Dutra et al., 2018; Wessel & Huber, 2019). Furthermore, the frontal cortex regions of right inferior frontal cortex and pre-supplementary motor area that are reliably active during successful response inhibition are likewise active when inhibition is required and following unexpected events that do not instruct response inhibition (e.g., Levy & Wagner, 2011; Sebastian et al., 2021). Together, these findings provide evidence for common inhibitory processes that are active during both action stopping and surprise processing.

A missing link in this picture concerns the role of β-frequency burst events, which have attracted considerable recent attention as a newly discovered inhibitory signature in the local field potential. β-bursts are transient, non-linear activity increases in β band activity (15-29 Hz) that occur in both cortical and subcortical regions of the brain (Feingold et al., 2015; Tinkhauser et al., 2018, Diesburg et al., 2021). β-bursts have been shown to represent time-frequency dynamics at the single-trial level more accurately and better predict behavior than average β power (Shin et al., 2017; Little et al., 2019; Wessel, 2020) and are potentially generated by a straightforward biophysical mechanism that maps onto inhibitory and excitatory thalamocortical dynamics (Sherman et al., 2016). In line with their purported role in reflecting inhibitory neural processes, several studies have recently linked β-bursts to motor inhibition. Little et al. (2019) showed that sensorimotor β-bursts that occurred close to movement increased RT. Wessel (2020) demonstrated that healthy adults exhibit decreased bilateral sensorimotor β-bursts in the moments leading up to motor execution along with further decreases in contralateral sites just before movement. The same study and subsequent work have identified relationships between β-bursts at more frontal sites and action cancellation. Wessel (2020) found that successful stop trials were characterized by an increase in frontocentral β-bursts which in turn prompted increased β-bursting at the bilateral sensorimotor sites. Jana et al. (2020) similarly observed an increase in frontal β-bursts on successful stop trials and found that their latency correlated with the latency of the downturn of subthreshold electromyographic activity evoked on a portion of successful stop trials. Because these increases in β-bursts occurred at around 120 ms following the stop signal and precede global motor suppression recorded TMS (∼140 ms), Jana et al. (2020) suggested that these β-bursts may be a signature of the triggering of the stop process (see also Choo et al., 2022). Diesburg et al. (2021) identified β-burst increases in local field potential recordings from subthalamic nucleus (STN) and motor thalamus on successful stop trials and further showed that these subcortical β-burst were associated with subsequent, low-latency sensorimotor β-bursts. Tinkhauser et al. (2018) found resting state phase coupling between STN and motor cortex that was increased during β-bursts. This suggests β-bursts may provide a vehicle of long-range communication during inhibitory motor control. Finally, β-bursts have been linked to pathophysiology of disease (Anidi et al., 2018; Lofredi et al., 2019; Pina-Fuentes et al., 2019), and clinical procedures aimed at targeting STN β-bursts in individuals with Parkinson’s Disease have shown promise in alleviating motor symptoms (Tinkhauser, Pogosyan, Little, et al., 2017; Tinkhauser, Pogosyan, Tan, et al., 2017).

Given the above-mentioned tight link between surprise and inhibition, and given the tight relationship between the novel β-burst signature and inhibitory control, one straightforward hypothesis is that β-bursts should be increased after unexpected events, even those that do not explicitly instruct the stopping of actions. Such a demonstration would be important for establishing a role of β-bursts in reactive inhibitory control. By reactive inhibitory control, we mean inhibition that is triggered by a stimulus rather than its anticipation (Aron, 2011). In the SST, the stop signal occurs on only a fraction of trials and uncertainty surrounds when it will occur. Consequently, some reactive inhibitory control is likely triggered by the stop signal. However, participants are not only aware that the stop signal will occur but are explicitly monitoring for it. Consequently, in the SST, reactive inhibitory control processes are necessarily conflated with proactive inhibitory control, or adjustments made in anticipation of the expected stop signal (Aron, 2011). Unexpected events offer a cleaner window into reactive control because such events prompt inhibition even when none is required by the task, and indeed even when inhibition may be antithetical to task goals (Wessel, 2018a). In addition, non-selective CSE suppression, which is a direct measurement of the physiological suppression of the motor system (Duque et al., 2017) is evident following unexpected events (Wessel et al., 2013; Iacullo et al., 2020). Such inhibitory control signatures that are observed both during instructed action-stopping (such as in the SST) and after unexpected events likely reflect reactive inhibitory control (though of course they may be modulated by the presence of proactive control, e.g., in the SST).

The primary goal of the current study was hence to investigate whether frontal β-bursts, known to be found after stop-signals, are also elevated following unexpected events, indicating their role in reactive inhibitory control. To this end, we used two datasets. In the first dataset, participants completed a trimodal cross-modal oddball task in which unexpected events could occur randomly and without warning in auditory, visual, and haptic modalities. In the second dataset, participants completed a bimodal cross-modal oddball task in which unexpected events occurred in either the auditory or visual modalities. This second dataset was previously reported by Wessel and Huber (2019) but is here newly analyzed for β-bursts. Regarding the first dataset, we hypothesized that frontal β-bursts would be increased in all three modalities. The second dataset then served as replication for the question of whether elevated β-bursts would be found following unexpected events. As additional research questions, we also examined the relationship of these β-bursts to surprise model estimates and RT at the single-trial level. To this end, we used the first dataset exploratorily and the second dataset (from Wessel & Huber, 2019) to independently verify (as initial confirmatory evidence) any exploratory findings generated from the first dataset.

## METHOD

### Participants

Dataset 1 includes 40 healthy young adult volunteers (21 female; 5 left-handed; *M*_Age_ = 21.35, *SD* = 4.17, range = 18 to 40). Data from three additional participants were excluded (two because of technical error in the CMO task and one who was unable to complete the CMO task). Dataset 2 includes the 55 healthy young adults from Wessel and Huber (2019). Participants in both datasets were recruited from the Iowa City community and from among University of Iowa students seeking research credit for psychology classes. Participants were compensated with course credit or at a rate of $15/hour. Both experiments were approved by the local institutional review board (#201511709).

Previous work on β-bursts from our group has shown that, in the stop-signal task (SST), increased FC β-bursts for successful stop compared to matched go trials show a large effect size (*d*z = .8). Subsequently, as few as 23 participants would be sufficient to attain 95% power for detecting an effect of equal size at an alpha level of .05. However, greater power is often required to establish reliable brain-behavior relationships. Prior work from our group on proactive β-bursts evinced such relationships with a sample of 41 participants (Soh et al., 2021). We therefore targeted a similar sample size in Dataset 1.

Prior to experimentation, piloting was performed to determine whether the unexpected events induced startle responses. A small group of participants completed the task while surface electromyography (EMG) was recorded from bipolar electrodes placed on the sternocleidomastoid. The continuous EMG was monitored, and visually detectable EMG bursts were interpreted as startle response. An initial, higher intensity vibration was found to produce startle response. We therefore opted for a lower intensity vibration experimentally and verified in one additional participant (after the main experiment) that none of the unexpected events in the CMO task produced startle response. We then repeated the CMO task with the high intensity vibration and verified that the high intensity vibration did produce the startle response.

### Materials

The methods used in Dataset 1 are the same as in Dataset 2 except where otherwise noted. The methods reported here are adapted from Wessel and Huber (2019). Stimuli for both experiments were delivered using Psychophysics toolbox 3 (Brainard, 1997) on MATLAB 2015 (TheMathWorks, Natick, MA) on an IBM-compatible computer running Fedora Linux. Visual stimuli were presented on an ASUS VG278Q low-latency flat screen monitor (144 Hz), and auditory stimuli were played at conversational levels (∼70 dB SPL) through speakers positioned on either side of the monitor. Tactile stimuli were delivered over custom-built response devices (Engineering Electronics Shop, University of Iowa, Iowa City, IA) that interfaced with a Psychtoolbox compatible data acquisition device (UBS-1208FS, Measurement Computing Corporation, Norton, MA) to record button presses from the thumbs and vibrate motors at rates of 8300 RPM. The response/vibration device were only used in Dataset 1. In Dataset 2 (and during the SST for participants in Dataset 1), participants made responses with each index finger on a QWERTY keyboard.

### Procedure

A diagram of the cross-modal oddball (CMO) tasks used in each dataset can be found in Fig. 1. In Dataset 1, each trial, started with a white fixation cross centered on a black background that was displayed for 500 ms. This was followed by the cue which was displayed for 200 ms. On 80% of trials, the cue was a standard event consisting of a green circle and a 600-Hz sine wave tone. On the remaining 20% of trials, 6.7% of trials featured unexpected visual events, 6.7% featured unexpected auditory events, and 6.7% featured unexpected haptic events. Thus, the unexpected events could occur in any (but not multiple) sensory domain(s). The specific trials on which unexpected events occurred were pseudo-randomly determined separately for each participant. The only stipulations were that the first three trials were always standard trials, equal numbers of each type of unexpected event were presented within a given block, and trials with unexpected events were always separated by at least one standard event trial. On unexpected visual trials, the green circle of the standard event was replaced by one of seven shapes (upwards triangle, downwards triangle, square, diamond, cross, hexagon, or serifed “I” shape) and one of 15 non-green colors equally spaced across the RGB spectrum. Thus, the unexpected visual events were also unique, or novel. Likewise, on unexpected auditory trials, the tone was replaced by a novel birdsong that matched the tone in amplitude. On unexpected haptic trials, the response devices that participants held in either hand vibrated bilaterally. The participants were instructed in advance about the properties of the standard event and that it was important to pay attention to this cue because it always took the same amount of time before the target would appear after the cue. Following presentation of the cue, the fixation was displayed for 300 ms prior to the presentation of the imperative stimuli. Thus, there was always a fixed delay of 500 ms before target onset. The target featured a left or right pointing arrow. The direction was pseudo-randomly determined for each participant with the only stipulation that each trial type contained an equal number of left and right facing arrows within each block. Depending on the direction the target arrow pointed, participants pressed a button on the response/vibration device with the thumb of their corresponding hand. Participants were instructed to respond as quickly and accurately as possible and had to respond within 1000 ms following the presentation of the target. After the response deadline, a variable inter-trial interval of 2600 to 3100 took place (in 100 ms steps, sampled from a uniform distribution) to preclude anticipation of the onset of the cue in the subsequent trial. Before beginning the formal experiment participants completed 10 practice trials with no unexpected events. They then completed 240 trials in the main experiment which were divided into 4 blocks of 60 trials with self-paced breaks in between.

**Fig. 1.**
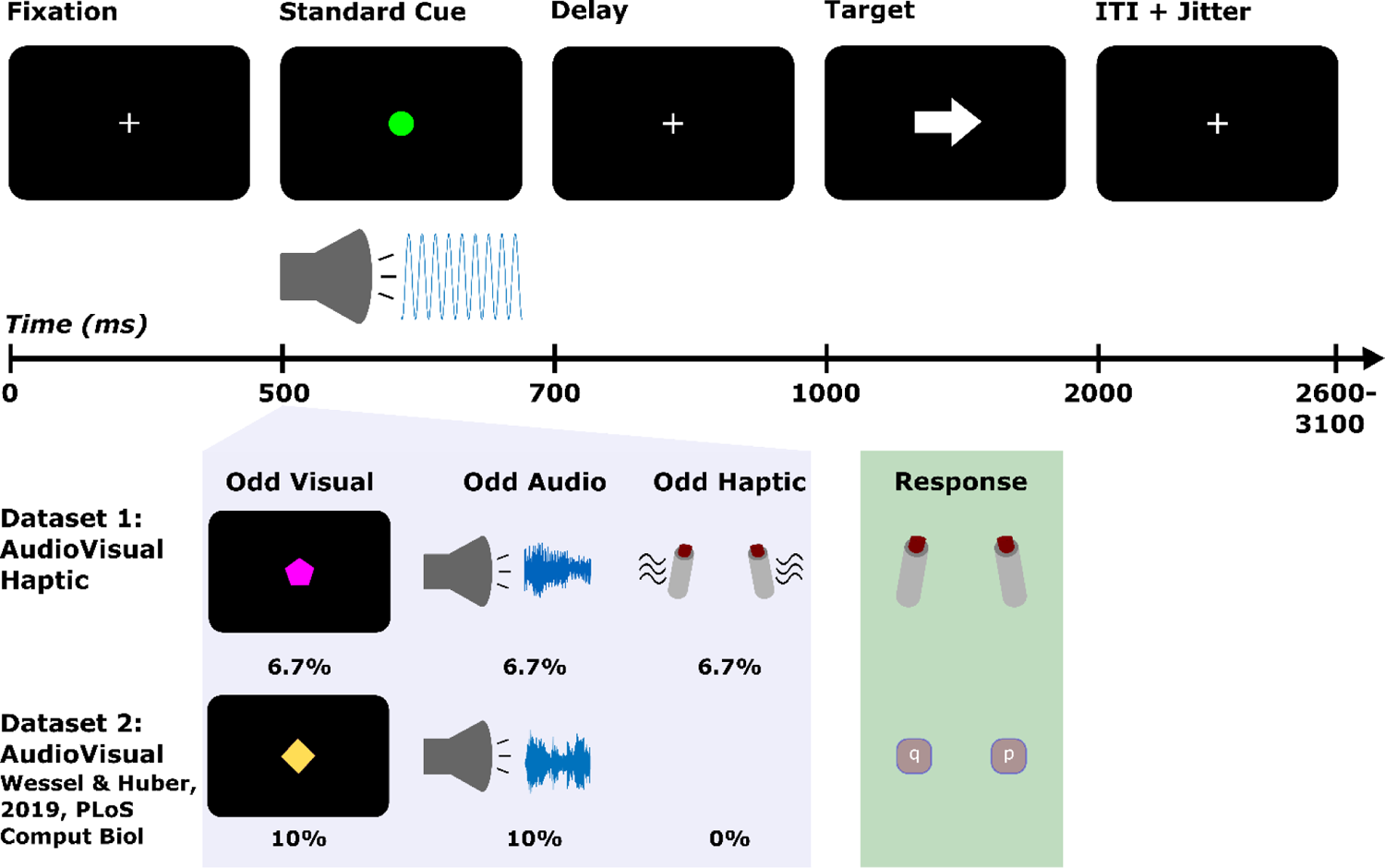
Task diagrams for the cross-modal oddball tasks in Datasets 1 and 2. The top row shows the timing and the display on standard cue trials (which were the same in both datasets). The standard cue consisted of a green circle and a 600 Hz pure tone. On trials with unexpected events, these were replaced by visual (unique shape/colors) or auditory novels (birdsong) or in the case of unexpected haptic events (in Dataset 1) the response devices participants held additionally vibrated. The light purple box illustrates examples of the unexpected events as well as their respective probabilities. The light green box depicts response devices (for Dataset 1, bilateral thumb presses, and for Dataset 2, bilateral index keypresses on a QWERTY keyboard).

The CMO task used in Dataset 2 was identical to Dataset 1 with two exceptions: In Dataset 2, 10% of trials featured unexpected auditory events and 10% featured unexpected visual events (thus, the overall frequency of the standard and unexpected events was kept the same across datasets 80/20%). Participants also responded by pressing buttons on a QWERTY keyboard with their index fingers (“q” with the left index for left-pointing targets and “p” with the right index for right-pointing targets).

Following the CMO task, participants in both datasets completed the SST. The SST was included to verify that the same inhibitory signatures of interest (frontal β-bursts, fronto-central P3) could also be identified following the Stop signal. To this end, the SST was used as a functional localizer to identify whether the same electrodes captured β-burst differences across tasks and whether the fronto-central P3 following unexpected events could be recovered from an independent component reflecting the fronto-central P3 following the Stop signal (as in Wessel and Huber, 2019). As in Dataset 2, participants completed a standard visual SST consisting of 300 trials completed in six, 50 trial blocks. The stop-signal probability was .33 resulting in 200 Go trials and 100 Stop trials. The trial began with a black fixation cross presented for 500 ms on gray background. After this, the left or right-pointing arrows (in black) were presented as the Go signal. On Go trials, participants pressed “z” key on a QWERTY keyboard in response to left arrows and the “m” key in response to right arrows. On Stop trials, the arrow turned red after the stop-signal delay (SSD) and participants were instructed to try their best to cancel their response while still responding as quickly and accurately as possible to the go signals. The SSD was initialized to 200 ms (separately for each hand) and increased in 50 ms increments following successful stop trials and decreased in 50 ms following failed stop trials. Trial duration was fixed at 3000 ms. Participants completed 24 practice (8 Stop-signal) trials before the formal SST experiment began. One person followed instructions to respond as quickly as possible for the CMO task but did not do so for the SST (as evidenced by SSDs of 1000 ms indicating extreme waiting for the stop signal and nullifying the ability of the task to measure action cancellation). Data from this individual were only excluded from analyses involving the SST data.

### EEG recording

Scalp-surface EEG was acquired from a 62-channel passive electrode cap connected to BrainVision MRplus amplifiers (BrainProducts, Garching, Germany). Additional electrodes were placed over the left canthus of the left eye and beneath the left eye (on the orbital bone). The ground electrode was located at the Fz electrode position, and the reference electrode was placed at the Pz electrode. EEG was digitized at a sampling rate of 500 Hz.

### EEG preprocessing

Preprocessing was performed as described in Wessel and Huber (2019). We used custom MATLAB (MATLAB 2015b, TheMathWorks) functions using the EEGLAB toolbox (Delorme & Makeig, 2004). The imported data were filtered with symmetric two-way least squares finite impulse response filters (high-pass cutoff: .3 Hz, low-pass cutoff: 30Hz). Outlier statistics were used to automatically reject non-stereotyped artifacts (joint probability and joint kurtosis with cutoffs set to 5 SD, cf., Delorme et al., 2007). The data were re-referenced to the common average and subjected to temporal infomax independent component analysis (ICA) decomposition (Bell & Sejnowski, 1995) with extension to sub-Gaussian sources (Lee et al., 1999). Among the resulting components, those corresponding to eye-movement and electrode artifacts were removed from the data based on outlier statistics and non-dipolar components with residual variances greater than 15% (Delorme et al., 2012).

### Behavioral analysis

The RT data were initially submitted to a Bayesian repeated-measures (RM)-ANOVA with the factors Event type (Standard, Unexpected audio, Unexpected haptic, and Unexpected visual) and Block (1, 2, 3, and 4) as within-subject factors. Moreover, as planned comparisons, Bayesian paired *t*-tests were used to compare RTs following each type of unexpected event to those following the standard event within each block (for a total of 12 comparisons). Error rate and miss rate were similarly analyzed with Bayesian RM-ANOVAs and Bayesian paired-tests. For these analyses and subsequent Bayesian tests, the data were analyzed using JASP 0.15.0.0 (Love et al., 2019) after exporting the data from MATLAB. Throughout, Bayes Factors (BF) are framed as evidence favoring the alternative hypothesis (H1) or the null hypothesis (H0) with *BF* ∼ 1 corresponding to inconclusive, *BF* > 1 corresponding to anecdotal evidence, *BF* > 3 corresponding to moderate evidence, and *BF* > 10 corresponding to strong evidence. For all tests, non-informative (or “empirical”) priors are used.

The subsequent SST functional localizer task was examined in terms of the mean Go-trial RT, mean failed-stop RT, and mean Stop-signal RT (estimated via the integration method, Verbruggen & Logan, 2009; Boehler et al., 2014).

### Modeling surprise at the single-trial level

There is some debate in the literature regarding the appropriate mathematical quantification of surprise (e.g., Shannon, 1948; Baldi & Itti, 2010; O’Reilly et al., 2013).

Based on Shannon’s theory of information, surprise is given by the following equation:

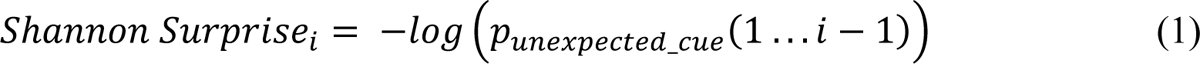

This equation quantifies the unexpectedness of the *i*th trial by taking the log inverse of the prior probability of the unexpected event. Since -log(0) is undefined, the surprise value for the first unexpected event (for which the prior is 0) was here defaulted to the next largest integer above the surprise value of the second unexpected event. The surprise values were different for each participant and computed independent of subject responses. Fig. 2 provides an example of the Shannon surprise values from one participant from Dataset 1.

**Fig. 2.**
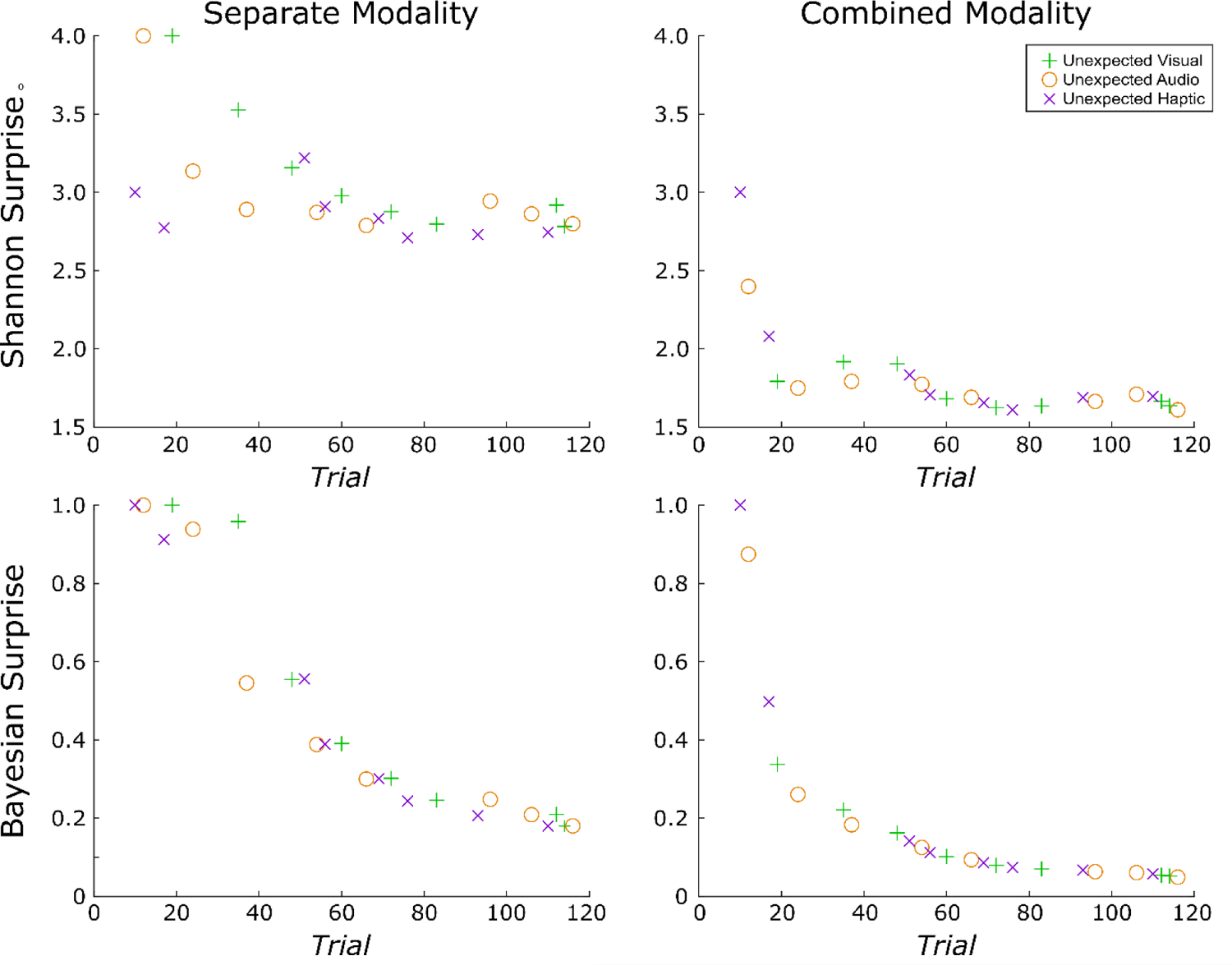
Example of the unique surprise model values from a single subject in Dataset 1. Surprise was quantified along two dimensions each with two levels. First, surprise was quantified according to each of two different formulas: (1) Shannon surprise, which is given by the inverse log of the probability of an unexpected event prior to its occurrence and (2) Bayesian surprise, which quantifies the difference in this probability and the updated probability after the unexpected event has occurred. Next, surprise was computed separately for each modality or combined (such that an unexpected haptic event is less surprising following an unexpected visual event). The ordering of trials containing unexpected events and thus the corresponding surprise values under each model varied across participants. Surprise values were computed independent of subject responses. While the rate of decay is similar for combined modality models calculated using either surprise term, Shannon and Bayesian surprise differ somewhat when calculated separately for each modality. Namely, the decay is less rapid for Shannon surprise and is more affected by variations in the number of trials between successive unexpected events of the same variety.

In contrast, Bayesian surprise (Baldi & Itti, 2010), an alternative quantification, uses the Kullback-Leibler divergence, and is given by the following equation:

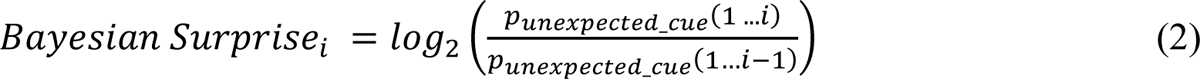

Bayesian surprise captures the difference between the prior (denominator) and posterior probabilities of an event (numerator). For this reason, some have argued that Bayesian surprise should be more accurately termed ‘model updating’ or ‘learning’ (e.g., O’Reilly et al., 2013; Nassar et al., 2019). Here, the first surprising value following the first unexpected event was defaulted to 1 because -log(0) (and division by zero) is undefined.

Notably, both Shannon and Bayesian surprise contain the prior probability of an event and are therefore correlated (though they can be potentially disentangled through experimental instructions, as in O’Reilly et al., 2013). An important distinction, though, is that whereas Bayesian surprise considers the difference in prior and posterior expectations of a given stimulus (based on its relative likelihood before and after a given event), Shannon surprise provides a scaled version of the prior expectation alone and does not take the posterior expectation into account. This creates important conceptual distinctions between the different surprise models. Most notably, if a substantial number of standard trials occur before a given unexpected event, Shannon surprise can increase relative to the preceding unexpected event. By contrast, Bayesian surprise is always strictly monotonically decreasing. In the current CMO tasks, Shannon surprise better represents local deviations in the unexpectedness of events whereas Bayesian surprise represents learning the expected probabilities themselves (Wessel and Huber, 2019). It is also worth noting that some have argued that Shannon surprise relates more to reorienting attention and effecting rapid action adjustments whereas Bayesian surprise operates at a higher order and relates more to updating beliefs or internal models about the environment (O’Reilly et al., 2013).

Beyond the method for quantifying surprise (Shannon vs. Bayesian), in a dataset that contains unexpected events originating in different modalities, a second question is whether surprise is modeled commonly across modalities or for each modality separately (Wessel & Huber, 2019). This consideration has a substantial impact on the resulting surprise values. In the combined model, the surprise values decay even more rapidly as an unexpected event in the auditory modality is deemed less surprising even if it follows an unexpected event in the visual modality. Wessel and Huber (2019) modeled Bayesian surprise for both common and separate models and found that frontocentral P3 amplitude was better predicted by the separate model, suggesting that frontal cortex separately accounts for these various sources of surprise. (For a replication and extension of this work with Dataset 1, please refer to the Appendix.) Of main interest, the present work provides a first examination of whether FC β-bursts relate to surprise. To this end, we again perform a model comparison that does not only compare Shannon vs. Bayesian surprise, but also compares separate vs. common surprise models across sensory modalities. Because the specific trials containing unexpected events and their ordering varied across participants, this also introduced variability in the corresponding surprise values under each model for each participant. Thus, any implicated surprise models in our single-trial model fitting analyses must be robust to this variability.

To complement the mean-based approach that examined RTs as function of Event Type and Block (detailed in the previous subsection), we also fit surprise values to RTs at the single-trial level. To do so, standardized (*z*-scored) surprise values were regressed onto standardized (*z*-scored) RTs using robust regression [in MATLAB, robustfit()] to generate beta weights for each participant. Group-level inference was drawn on the resulting beta weights using Bayesian one-sample Wilcoxon-signed rank tests to assess whether the surprise values reliably fit the data.

### β-Burst extraction

Before identifying β-bursts, we minimized the potential for volume conduction by transforming the data to a reference-free montage via the current-source density (CSD) method (Perrin et al., 1989; Tenke & Kayser, 2005).

β-bursts were identified in the same manner as described by Shin et al. (2017) and Wessel (2020). The only exception is that we used a 2× median power threshold (in lieu of a 6× median power threshold) based on Enz et al.’s (2021) specification of optimal settings for β-burst detection. The description is adapted from Shin et al. (2017) as implemented in Wessel (2020):

First, each electrode’s data were convolved with a complex Morlet wavelet of the form:

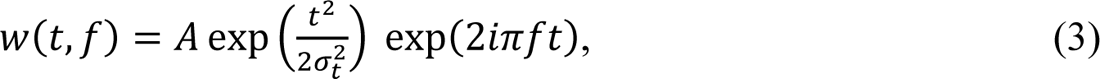

with 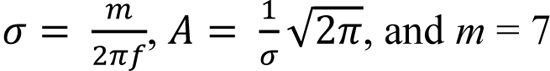 (cycles) for each of 15 evenly spaced frequencies spanning the β-band (15–29 Hz). Time–frequency power estimates were extracted by calculating the squared magnitude of the complex wavelet-convolved data. These power estimates were then epoched relative to the events in question (ranging from −500 to +2000 ms with respect to the cue in the CMO task, and −500 to +1000 ms with respect to signal onset in the SST). Individual β-bursts were defined as local maxima in the trial-by-trial β-band time–frequency power matrix for which the power exceeded a set cutoff of 2× the median power (Enz et al., 2021) of the entire time–frequency epoch power matrix for that electrode (i.e., the median thresholding was performed across all epochs). Local maxima within each epoch were identified using the MATLAB function imregionalmax().

### β-Burst analyses in the CMO tasks

We focused on a frontocentral region of interest for the β-burst analysis based on our previously reported finding involving a large SST dataset (n=234) showing maximal differences in beta burst rate in these electrodes on successful Stop compared to match Go trials (Wessel, 2020). To examine whether FC β-bursts were elevated following unexpected events and to potentially gain a better understanding of the time course of the β-bursts, we adopted a sliding window approach in place of the bin-based approaches used in prior work (e.g., Wessel, 2020). For this, we cycled through each sample beginning −100 ms prior to the onset of the cue and +500 ms after its onset and counted the number of bursts occurring within +/- 25 ms of the sample (with one sample corresponding to 2 ms of data). We then converted to burst rate by summing the number of bursts across trials and dividing by the number of trials. This process was completed for each of the FC electrodes of interest (FC1, FCz, FC2, C1, Cz, and C2). After this, we used the baseline period (−100 to 0 ms) to calculate the percent change in burst rate and to convert to percent change from baseline for each electrode. The resulting values were averaged across the FC electrodes. For these and all subsequent analyses, standard event trials that were immediately preceded by unexpected event trials were excluded. The data were then compared across each Event Type (standard, unexpected audio, unexpected visual, unexpected haptic). First, we used frequentist one-way repeated-measure ANOVAs over each time point comparing percent change in burst rate across the various event types. Then, we used frequentist paired samples *t*-tests to compare each unexpected event type to the standard event. The initial, pre-defined alpha-level of .05 was used for these tests. The alpha-level was then adjusted for multiple comparisons via the FDR procedure (Benjamini et al., 2006) correcting for the number of tests (one for the ANOVA, three for the paired *t*-tests in Dataset 1, and two for the paired *t*-tests in Dataset 2) and the number of timepoints (250).

Next, the relationship between β-burst rate and surprise was explored. In each dataset, we looked at the association between the number of β-bursts and surprise values on (non-rejected) trials containing unexpected events. In our initial, exploratory analysis with Dataset 1, we considered two different time ranges. First, we considered beta-bursts within the full cue-to-target interval (0 to +500 ms). As these results were exploratory and did not yield positive findings, we do not consider them further. Second, and after having established that, in line with our main hypothesis, FC beta-bursts were elevated following unexpected events in Dataset 1, we explored these relationships within a specific time window containing the elevated β-bursts. For this, we extracted the number of β-bursts between +75 and +200 ms following each unexpected event. Both the number of bursts within these search windows and the surprise values (again for each type of surprise and for separate and combined modality considerations) were standardized as *z*-scores and a beta weight was computed using robust regression (with the robustfit() function in MATLAB) for each subject. To draw inferences at the group-level, the resulting individual beta-weights were compared against zero using Bayesian one-sample Wilcoxon signed-rank tests. We here consider the results of Dataset 1 to be purely exploratory, whereas we consider any potentially replicated results in Dataset 2 to provide initial, confirmatory evidence.

We then examined the relationship between RTs and β-bursts in each dataset using similar single-trial level approaches. First, we considered whether the number of β-bursts predicted RT by computing spearman rank correlations for each participant following unexpected events. For these analyses, we considered the specific time range (+75 to +200 ms following the event) and the full cue-to-target interval (0 to +500 ms) in both datasets. We then considered whether the latency of the β-burst in relation to the target was predictive of RT. As one approach, we extracted the latency of the β-burst that was nearest to the target (in the −250 to+250 ms surrounding the target) and again computed spearman rank correlations for each participant. As another approach, we compared RTs across trials that contained β-bursts in the FCz electrode (which was selected as a representative electrode based on results from the main analyses) in the first half of the cue-to-target interval, the second half, both, and neither.

### β-Burst analysis in the SST

Lastly, we examined the dynamics of FC β-bursts in the SST (combined across Datasets 1 and 2) using the sliding window approach from the CMO task to compare β-bursts on successful Stop trials to matched Go trials as well as to failed Stop trials.

## RESULTS

### Behavior

The RT results of Dataset 1 closely matched those previously reported for Dataset 2 (Wessel & Huber, 2019) in all respects. The Bayesian RM-ANOVA on Event Type (standard, unexpected audio, unexpected visual, unexpected haptic) and Block (1, 2, 3, 4) indicated strong evidence for main effects of Event Type (*BF_10_* = 8.02 x 10^4^) and of Block (*BF_10_* = 2.71 x 10^9^). Strong evidence was also indicated for the Event Type X Block interaction over a model with only the two main effects (*BF_10_* = 268.1). Most importantly, the basic comparisons between each type of unexpected event and standard events echoed the finding that all types of unexpected events induce RT slowing, but that — consistent with the notion of surprise adaptation — these differences wane over time. This was apparent as unexpected visual and unexpected auditory events both showed strong evidence of slowing compared to standard events in the first block (Unexpected audio: *BF_10_* = 53.6; Unexpected visual: *BF_10_* = 2.12 x 10^5^). Likewise, we found that Unexpected haptic events also showed this slowing in the first block (*BF_10_* = 5.72). Only anecdotal evidence was found for RT slowing following unexpected visual events in Block 2 (*BF_10_* =1.10) and Block 3 (*BF_10_* =2.64). None of the other unexpected events in Block 2 and onwards showed evidence of RT slowing (*BFs_01_* > 1). Thereby, the behavioral RT results replicated Wessel and Huber (2019) and extended them to the haptic domain.

The single-trial fits of RT and surprise lent themselves to similar conclusions as the mean-based approach to RTs, with some additional insights. There was strong evidence to suggest that Shannon surprise with separate modality terms positively predicted RTs (*BF_10_* = 20.49), whereas there was moderate evidence against Shannon surprise with common modality terms predicting the data (*BF_01_* = 4.48). Similarly, there was strong evidence that Bayesian surprise with separate modality terms positively predicted RTs (*BF_10_* = 24.26), but only anecdotal evidence to suggest this for Bayesian surprise with common modality terms (*BF_10_* = 2.18). Thus, participants’ RTs appeared to follow both types of surprise, provided their values represented surprise separately for each type of unexpected event.

For mean error rates, there was a main effect of Event Type on error rate (strong evidence, *BF_10_ =* 12.69) but not of Block (strong evidence, *BF_01_* = 10.99), nor was there an Event Type x Block interaction (strong evidence, *BF_01_* = 45.5). The mean percent error rates for each Event Type were as follows: Standard: 1.97%, Unexpected auditory: 0.16%, Unexpected visual: 1.09%, and Unexpected haptic: 1.41%. Follow-up *t*-tests on Event Type indicated that the main effect was driven by the lower number of errors following unexpected auditory events compared to standard events (*BF_10_* = 9.76 x 10^5^). The evidence did not suggest that the other types of unexpected events differed from the standard (unexpected visual: *BF_01_* = 2.42, unexpected haptic: *BF_01_* = 6.17), nor did they differ from the unexpected auditory event (unexpected visual: *BF_01_* = 1.22, unexpected haptic: *BF_01_* = 9.90). This outcome contrasts those for Dataset 2 which showed increased error rates following unexpected auditory and visual events that remained stable across blocks. However, error and miss trials were generally rare and not considered in the EEG analyses.

For mean miss rates, the results were in line with the null findings of Dataset 2. Strong evidence was found for a lack of differences regarding Event Type (*BF_01_* = 125.0), Block (*BF_01_* = 11.23), and the Event Type x Block interaction (*BF_01_* = 311.6).

The SST also closely followed those from Dataset 2. Mean Go RT = 535 ms, mean failed Stop RT = 464 ms, mean SSRTi = 252 ms, and mean *p*(inhibit) = .52 (compared to Dataset 2: mean Go RT = 520 ms, mean failed Stop RT = 444 ms, mean SSRTi = 252.00, and *p*(inhibit) = .51).

### EEG

#### FC β-bursts are increased following unexpected events in all three modalities

In Dataset 1, significantly increased β-bursting followed each type of unexpected event (see Fig. 4). The one-way RM-ANOVAs did not return significant differences after FDR-correction (sustained differences at p < .05 were found from 78-138 ms and at 212 ms without correction). However, more importantly, comparing the sliding window trace for each unexpected event to the standard revealed significant β-burst increases following unexpected auditory events from 102-148 ms, following unexpected visual event from 116-118 ms and following unexpected haptic events from 52-130 ms.

**Fig. 3.**
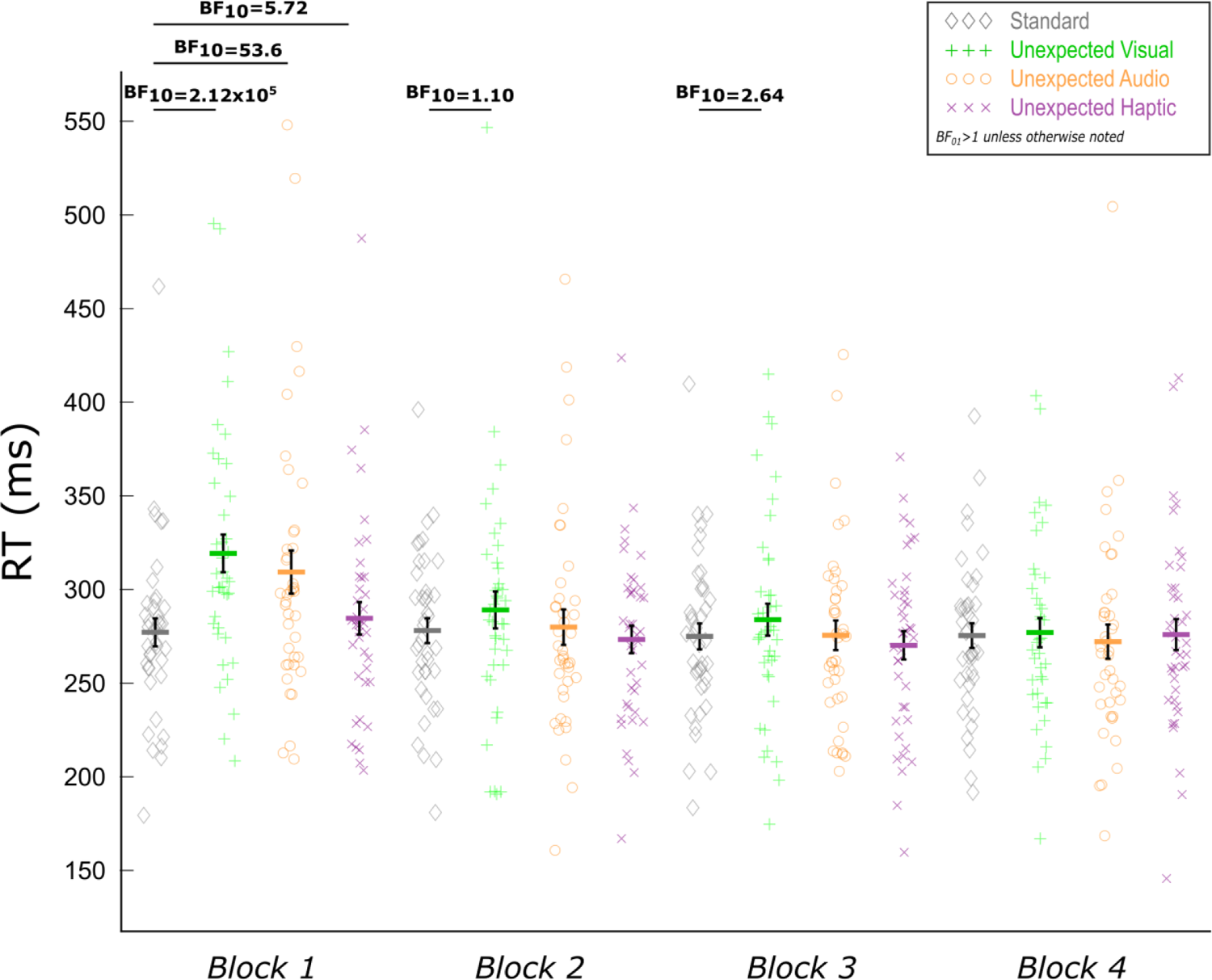
Mean RTs as a function of Event Type and Block for the trimodal cross-modal oddball task of Dataset 1. Paired Bayesian t-tests indicated RT slowing following all three types of unexpected events in the first block. Consistent with the notion of surprise, these differences quickly dissipated as suggested by the lack of differences in subsequent blocks (and the attenuated difference following unexpected visual events in the second block). Black error bars indicate SEM.

**Fig. 4.**
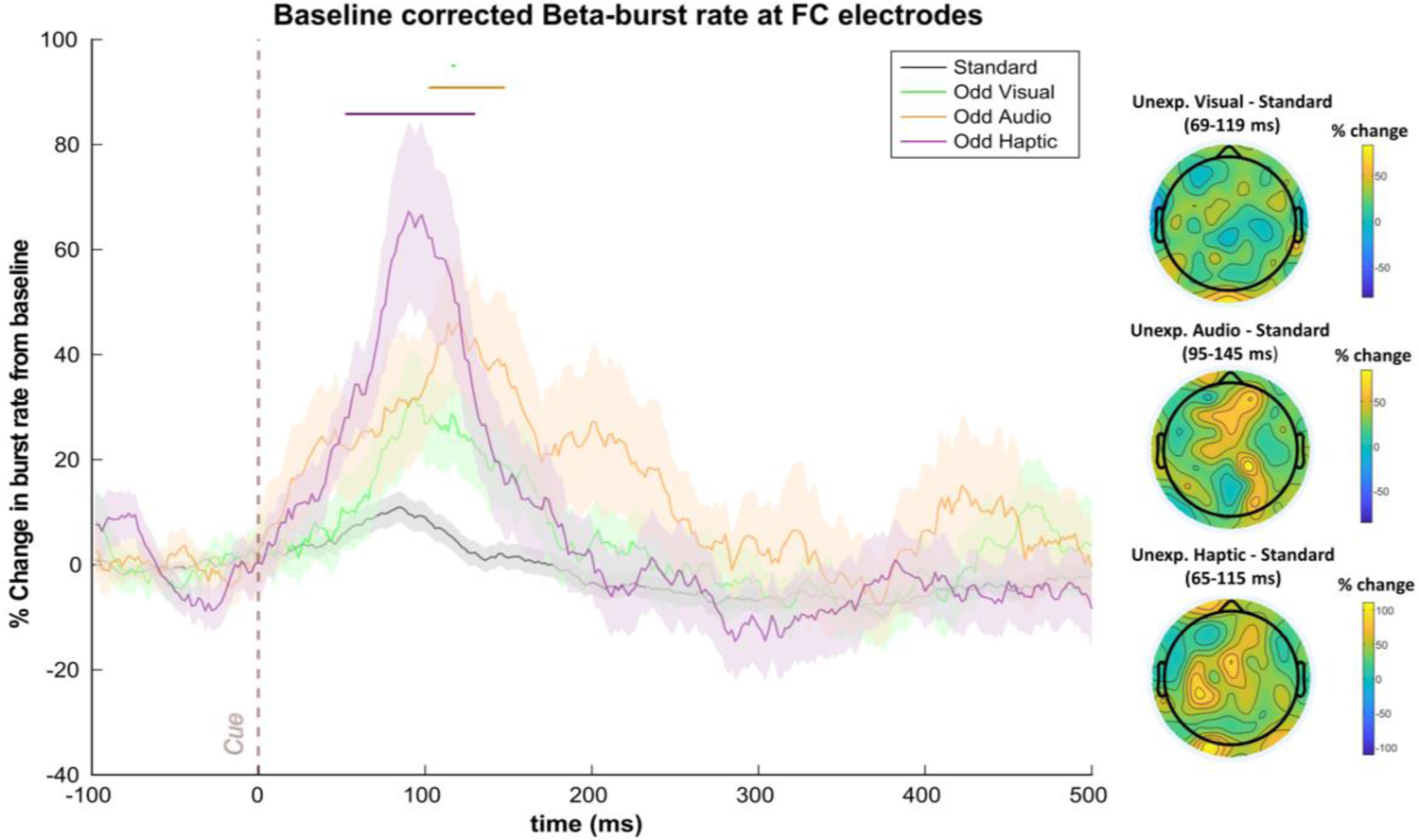
FC β-bursts following unexpected events in Dataset 1 (n = 40). (Left) Depicts the percent change in β-burst rate relative to baseline for +/- 25 ms sliding windows calculated for each 2 ms timepoint. An increase in β-bursts at FC sites was evident following unexpected events in all three modalities at early latencies. Sliding window timepoints with significant differences (FDR-corrected, p = .05) across event types are denoted by horizontal lines at the top of the plot (black: rm-ANOVAs across all event types, green: unexpected visual vs. standard, orange: unexpected auditory vs. standard; purple: unexpected haptic vs. standard). Shaded regions indicate SEM. (Right) Scalp topography differences depicting the % change in β-bursts at all electrodes in the +/- 25 ms window surrounding the maxima for each unexpected event type identified with the sliding window technique relative to the same time period for the standard event.

As can be seen in Fig. 4, the unexpected haptic events appeared to prompt more robust increases in β-bursts than unexpected events in the other sensory modalities. It is important to note, though, that differences other than the modalities in question could account for these differences particularly because the unexpected haptic events always featured the same stimulus (rather than a novel) and because no haptic event was not a part of the standard cue. Presumably, the lack of novelty among unexpected haptic events might have rendered them less unexpected whereas only encountering a haptic event on unexpected haptic event trials might have rendered such events more unexpected. Of course, we also cannot discount the possibility that fundamental differences between sensory modalities might also account for differences in β-burst characteristics following unexpected events. For instance, β power appears to underly communication between parallel sensorimotor representations of somatosensory and motor cortex (Baker, 2007) and sensorimotor β-bursts have been shown to play an important role in somatosensory perception (Shin et al., 2017; Law et al., 2022). Thus, β-bursts may be particularly important in the haptic domain. At present, we refrain from drawing comparisons across different types of unexpected events and instead focus on FC β-bursts that were common across different unexpected events and may thus reflect domain-general control mechanisms.

In Dataset 2, the RM-ANOVA detected significant differences across conditions from 110-232 ms (see Fig. 5). These sliding windows coincided virtually identically with those identified for the unexpected auditory events compared to the standard event (106-232 ms). Although unlike in Dataset 1, we did not find significant increases following unexpected visual events by themselves, this was a consequence of the FDR correction. Without it, (i.e., using p < .05), β-bursts were again increased following unexpected visual events (12-50 ms). This deviation from Dataset 1 notwithstanding, the overall results strongly suggest that, as predicted, β-bursts are generally increased following unexpected events.

**Fig. 5.**
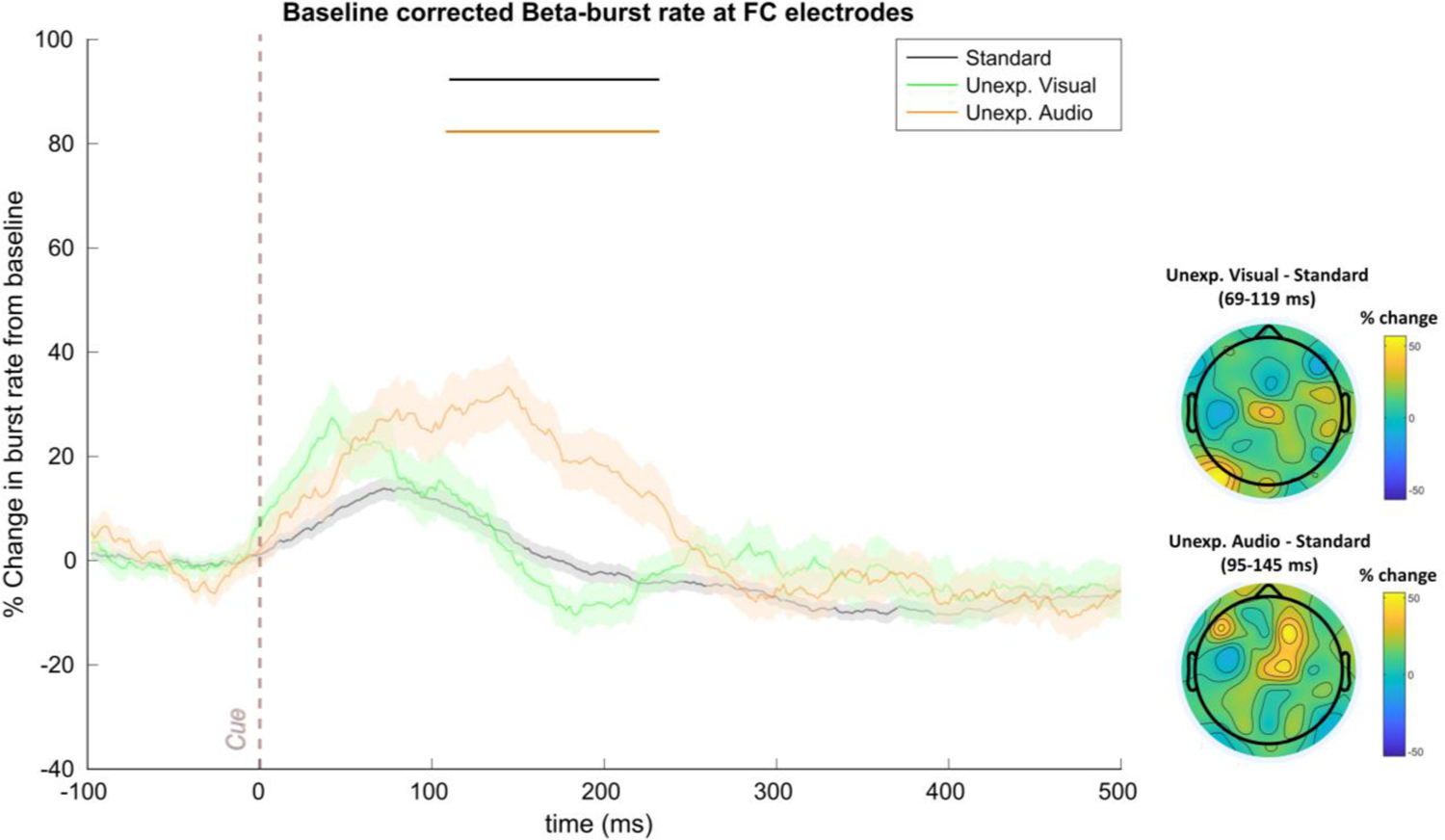
FC β-bursts following unexpected events in Dataset 2 (n=55). (Left) Depicts the percent change in β-burst rate relative to baseline for +/- 25 ms sliding windows calculated for each 2 ms timepoint. An increase in β-bursts at FC sites was evident following unexpected auditory events. The numerical increase in β-burst rate following unexpected visual events did not achieve significance. Sliding window timepoints with significant differences (FDR-corrected, p = .05) across event types are denoted by horizontal lines at the top of the plot (black: rm-ANOVAs across all event types, green: unexpected visual vs. standard, orange: unexpected auditory vs. standard). Shaded regions indicate SEM. (Right) Scalp topography differences depicting the % change in β-bursts at all electrodes in the +/- 25 ms window surrounding the maxima for each unexpected event type identified with the sliding window technique relative to the same time period for the standard.

As an alternative approach suggested to us by an anonymous reviewer, we repeated the above approach but, instead of using the pre-specified FC ROI, we used a cluster of frontal electrodes that was identified based on the corresponding SST that participants completed after the CMO task in each dataset. Specifically, we identified a group of contiguous frontal electrodes displaying significant (p < .001) increases in β-bursts for at least 10 consecutive time windows on Successful Stop compared to Go trials in the pre-SSRT period. Of the electrode clusters meeting these criteria, we selected the one showing the maximal difference. For Dataset 1, this cluster consisted of electrodes FCz, FC1, Cz, and C1. This analysis yielded highly similar results to the original analysis. Significant (FDR-corrected according to the original analysis) β-burst increases were found following unexpected auditory events from 100-150 ms; unexpected haptic events from 90-124, 128-132 ms; and unexpected visual events at 98, 102-112, and 116-118 ms. For Dataset 2, a cluster of electrodes consisting of FC1, FCz, FC2, C1, AF4, FC4, F2, F4, and F6 were identified as reflecting heightened β-bursts on successful Stop compared to Go trials. Significant β-burst increases were found following unexpected auditory events from 52-56 ms, 106-244 ms, and 254-272 ms; as well as following unexpected visual events from 58-62 ms and at 364 ms. These analyses thus complemented the original analyses in implicating early latency increases in β-burst rate following each type of unexpected event.

#### Early FC β-bursts reflect modality-specific Shannon surprise terms

We used Dataset 1 for exploratory analyses of potential relationships between FC β-bursts and surprise. We then aimed to replicate any such findings using Dataset 2. After having identified that FC β-bursts were increased following unexpected events in Dataset 1, we considered these relationships within a specific time range that contained the period during which unexpected events were followed by significant increases in β-bursts (75 to 200 ms after event onset). The individual-subject beta-weights from these analyses are plotted in Fig. 6a. Shannon surprise with separate modality terms for each modality was positively associated with β-bursting as *BF* indicated moderate evidence that the mean beta weight at the group-level was above zero (*BF_10_* = 7.20). Contrastingly, *BFs* for the other surprise models indicated moderate evidence for no relationship (Shannon surprise/combined terms: *BF_01_* = 5.81, Bayesian surprise/separate terms: *BF_01_* = 3.82, Bayesian surprise/combined terms: *BF_01_* = 2.40).

**Fig. 6.**
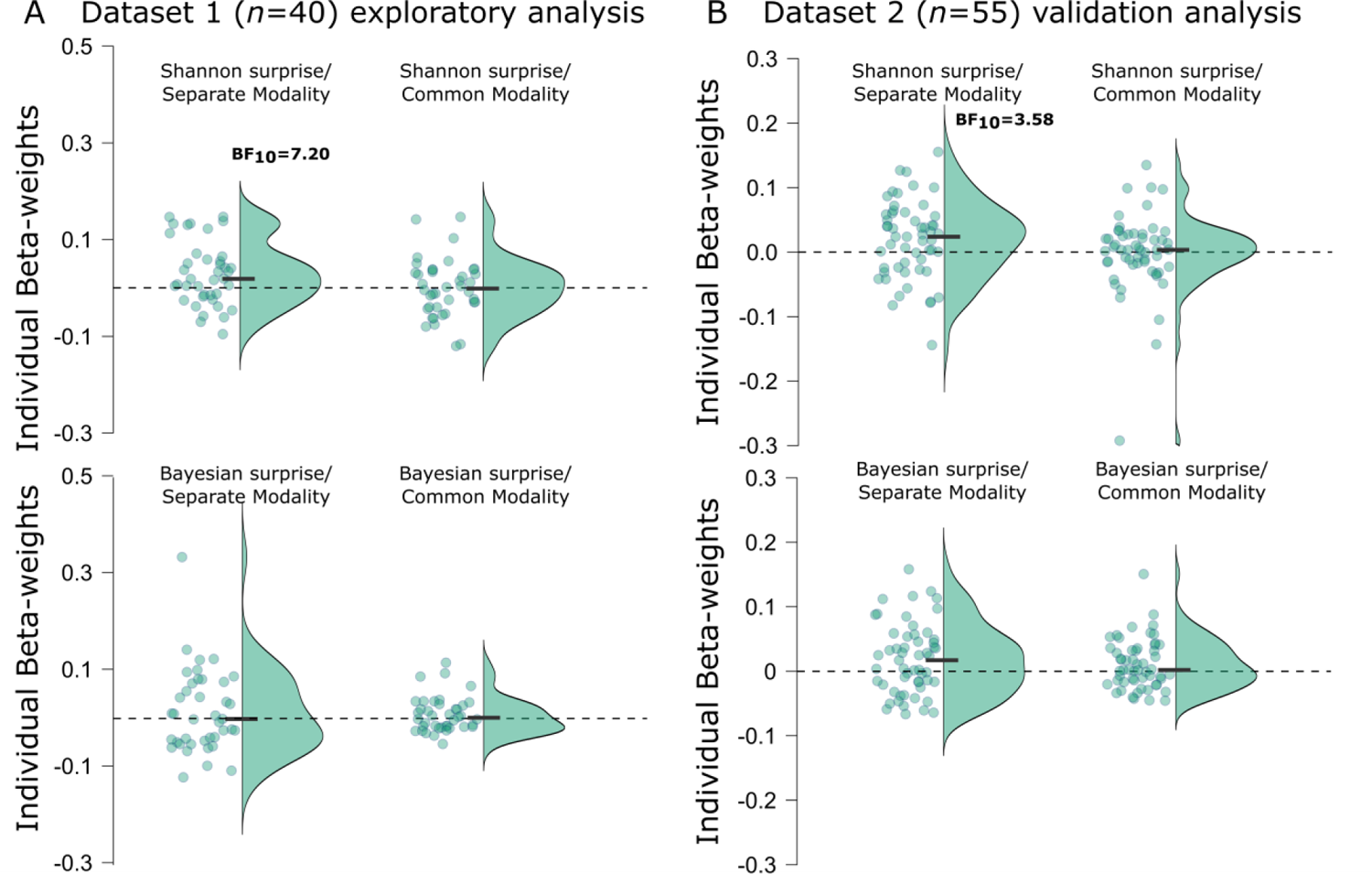
Individual beta-weights from the various z-scored surprise single-trial model values fitted to z-scored β-bursts from 75 - 200 ms following unexpected events. **a** Points represent mean beta-weights from individual subjects estimated via robust regression for each surprise model in Dataset 1. Half violin plots depict the distribution of beta-weights. Black bars indicate mean beta-weights. **b** The same as in panel a but for Dataset 2 which was used to validate potential findings in Dataset 1. Reported BF_10_s correspond to moderate evidence in favor of the alternative hypothesis. BF_10_s < 3 are not shown here (but are reported in the main text).

To confirm these exploratory findings, we completed the same analyses using Dataset 2. The individual-subject beta-weights are plotted in Fig. 6b. Just as in Dataset 1, there was moderate evidence that Shannon surprise with separate model terms positively predicted FC β-bursting (*BF_10_* = 3.58). While there was again moderate evidence for no relationship regarding Shannon surprise with combined modality terms (*BF_01_* = 6.54), this time, there was anecdotal evidence suggesting positive relationships for Bayesian surprise with separate terms (*BF_10_* = 2.53) and Bayesian surprise with combined terms (*BF_10_* = 1.29). Here, it may be worth mentioning that in the trimodal dataset, any given unexpected event was less frequent than in the bimodal dataset. Consequently, the difference between Shannon surprise and Bayesian surprise values should be relatively more pronounced in Dataset 1 owing to the longer stretches of trials between successive unexpected events of the same ilk. This could potentially explain why Bayesian surprise models appeared to show some hint of a relationship in Dataset 2 that was not observed in Dataset 1. Either way, Shannon surprise with modality-specific terms consistently outperforms the other models across both datasets.

We next sought to corroborate these findings using the ROIs based on the frontal electrodes showing increased β-bursts on successful stop trials in the SST participants completed after their respective CMO tasks. The specific electrodes used for each Dataset in these analyses are detailed in the previous section. For Dataset 1, there was strong evidence *BF_10_* = 400.01 indicating that Shannon surprise with separate modality terms positively predicted β-bursts. For the other surprise models, anecdotal to moderate evidence was provided against an association between surprise values and β-bursts (Shannon surprise/common modality: *BF_01_* = 5.81, Bayesian surprise/separate modality: *BF_01_* = 2.02, Bayesian surprise/common modality: *BF_01_* = 1.64). For Dataset 2, there was again strong evidence *BF_10_* = 35.01 indicating a positive association between Shannon surprise with separate modality terms and β-bursts. As with the original ROI, there was moderate evidence against an association for Shannon surprise with common modality terms (*BF_01_* = 3.16), anecdotal evidence for an association with Bayesian surprise with separate modality terms (*BF_10_* = 1.42), and anecdotal evidence for an association with Bayesian surprise with common modality terms (*BF_10_* = 2.35). Thus, these results also clearly establish a relationship between β-bursts and Shannon surprise with separate modality terms, although here, the β-bursts may bear a tighter connection to action stopping given that these participants showed increased β-bursting in the same electrodes during successful stopping in the SST.

#### No observed relationships between FC β-bursts and RT in the CMO task

We next investigated relationships between RT and FC β-bursts. No reliable associations were found between the number of FC β-bursts and RTs on unexpected event trials in either dataset with either time range (Dataset 1, full time range: *BF_01_* = 5.62, constrained time range: *BF_01_* = 2.67; Dataset 2, full time range: *BF_01_* = 5.41, constrained time range: *BF_01_ =* 6.67). Neither were reliable associations found on standard event trials (Dataset 1, full time range: *BF_10_* = 5.29, constrained time range: *BF_01_* = 5.68; Dataset 2, full time range: *BF_01_* = 6.76, constrained time range: *BF_01_* = 4.02). We also did not observe evidence for relationships between the latency of β-bursts in the −250 to +250 ms surrounding the onset of the target and RT (Dataset 1: Unexpected events: *BF_01_* = 4.85, Standard events: *BF_01_* = 4.74; Dataset 2: Unexpected events: *BF_01_* = 4.83, Standard events: *BF_01_* = 6.06). Finally, we compared RTs on trials depending on whether β-bursts in the FCz electrode (as a representative electrode across all ROIs) were found in the first half of the cue-to-target interval, the second half, both, or neither. Bayesian RM-ANOVAs did not indicate any differences among these conditions (Dataset 1: *BF_01_* = 16.13; Dataset 2: *BF_10_* = 1.27). In the Discussion section, we discuss potential reasons for the present lack of relationships between β-bursts and RT.

#### Stop-signal related FC β-bursts are increased later than those after unexpected events

As can be seen in Fig. 7, FC β-bursts were also increased following the Stop signal. In the SST (from Datasets 1 and 2, *n* = 94), FC β-bursts were increased on successful Stop trials compared to matched Go trials from 174-272 ms. This replicates prior work documenting increased β-bursting at frontal sites following the stop signal (Wessel, 2020; Jana et al., 2020). Interestingly, no time point differences were found for successful compared to failed stop trials. However, the peak burst rate latency for successful stops appeared to precede that for failed stops (Fig. 7). This parallels the main finding involving FC P3 in which the amplitude peaks earlier on successful Stop trials (e.g., Kok et al., 2004) and would suggest that the process related to FC β-bursts begins (or at least comes to fruition) sooner on these trials.

**Fig. 7.**
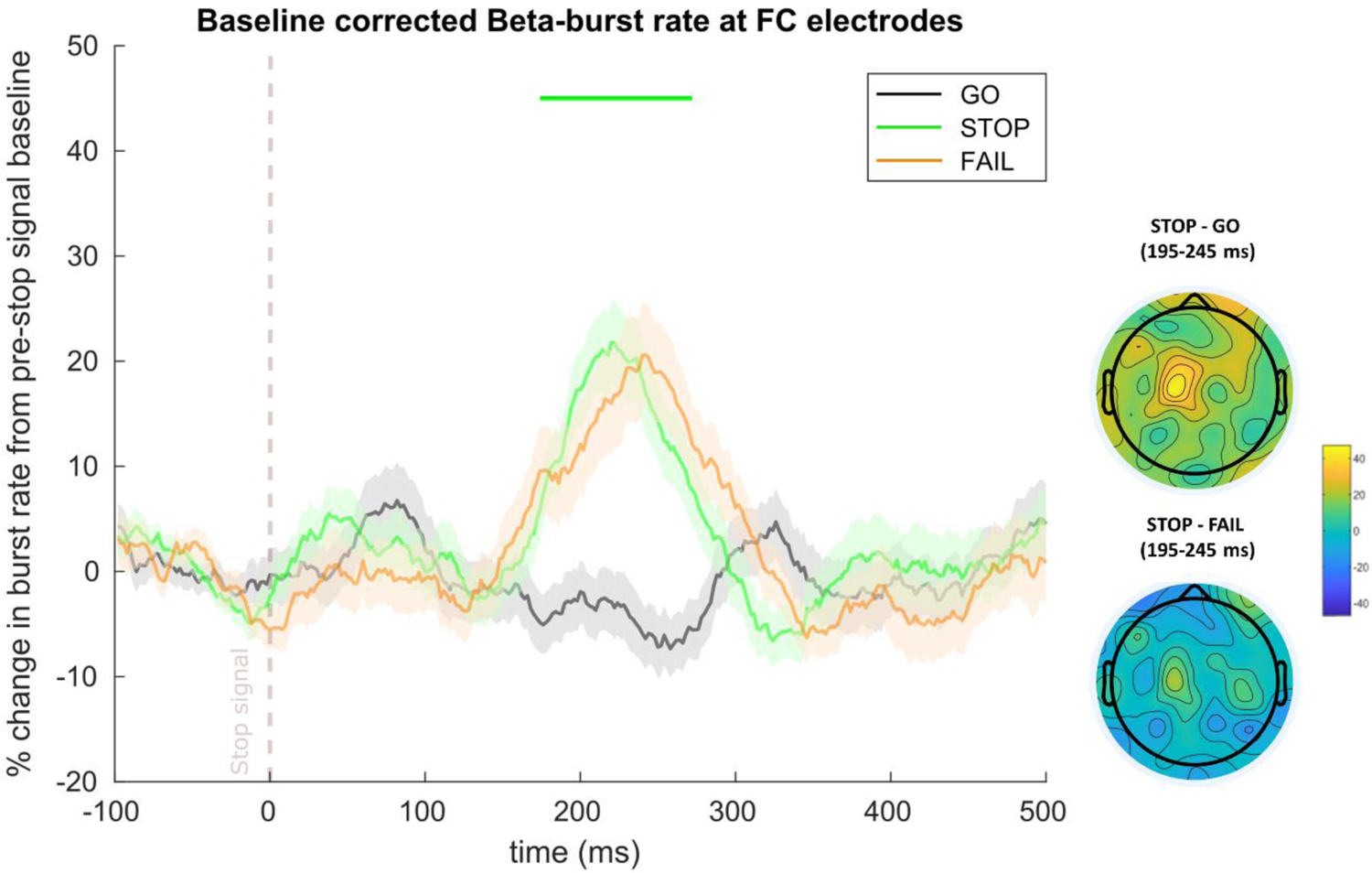
FC β-bursts following the stop signal in the SST (n=94, combined from Datasets 1 and 2). Depicts the percent change in β-burst rate relative to baseline for +/- 25 ms sliding windows calculated for each 2 ms timepoint. β-bursts at FC sites were increased on successful stop trials (stop) compared to matched Go (go). Failed Stop trials (fail) did not differ from successful Stop trials. Sliding window timepoints with significant differences (after FDR correction) across trial types are denoted by horizontal lines at the top of the plot (green: successful stop vs. matched go). Shaded regions indicate SEM. (Lower Right) Scalp topography differences depicting the % change in β-bursts at all electrodes in the +/- 25 ms window surrounding the maximum for successful stop compared to matched timepoints on go and failed-stop trials.

#### Timing differences of FC β-bursts and FC P3 following unexpected events in the CMO task and the Stop signal in the SST

Fig. 8 depicts the differences in FC β-burst rate following each type of unexpected event and the standard in the CMO task of Dataset 1 as well as the difference between successful Stop and matched Go trials in the corresponding SST. It is interesting that the time range of increased FC β-bursts following the stop signal appeared to occur later than the time ranges in which we found increased FC β-bursting following unexpected events in the CMO task. Similarly, an examination of the activity corresponding to the FC P3 as captured by the IC that best reflected the Stop signal P3 (i.e., that showed a maximum difference in activity at frontal sites and highest correlation to the original channel space ERP, Wessel & Huber, 2019) suggested that the P3 process appears to begin sooner following unexpected events in the CMO task than on successful Stop trials in the SST. Finally, the FC β-burst peaks in each condition and task also appeared to precede the earliest differences in the corresponding FC P3 (as isolated with the IC that best reflected each participant’s Stop signal P3).

**Fig. 8.**
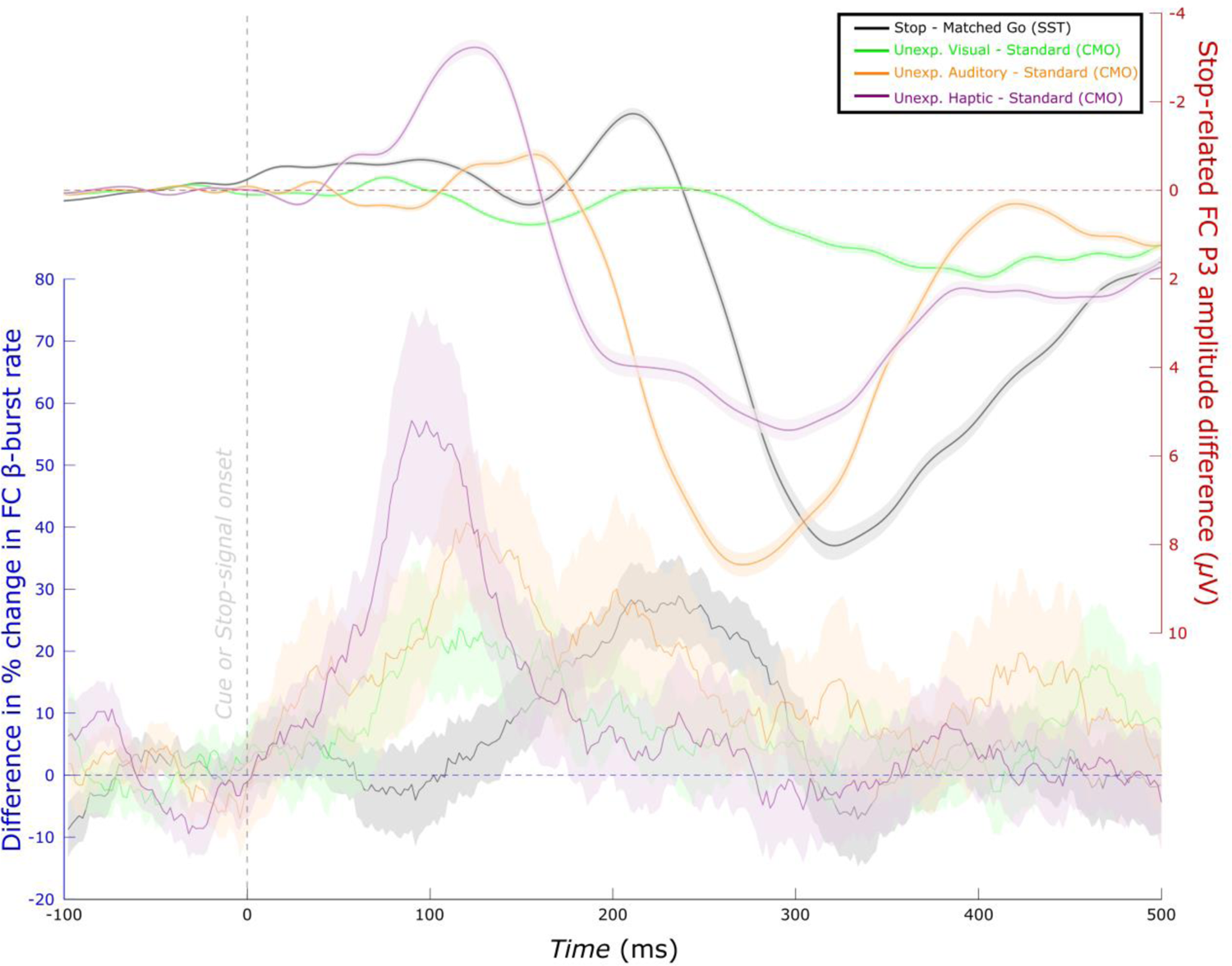
Time-course of differences in FC β-burst rate compared to FC P3 (Dataset 1, n=39). (Left Scale) Differences in the percent change in β-burst rate for successful Stop trials compared to matched Go trials (from Fig. 7, but only involving Dataset 1 participants) as well as unexpected events relative to standard (from Fig. 4). (Right Scale) The same differences for FC P3. For this, the SST was used as a functional localizer to isolate an IC corresponding to the FC stop-signal P3 in the merged dataset (i.e., the CMO and SST in Dataset 1). Visual inspection indicates that FC β-burst differences precede the corresponding FC P3 differences. Unexpected event differences generally also precede stop differences (excepting the FC P3 for unexpected visual events).

Following a suggestion from an anonymous reviewer, we completed an exploratory analysis to assess for statistical differences in these latencies at the individual-subject level. To examine latency differences in β-bursts, we used the sliding window approach for each participant to extract the peak burst rate within +50 to +400 ms following each type of unexpected event in the CMO task and following the Stop signal on successful Stop trials in the SST. To identify P3 onset, we first identified the maximum difference in the Stop signal IC in each participant during the same time range (although P3 is typically extracted with a later time range, here it was important to search within the same time range). We then used permutation testing (with Markov Chain Monte Carlo simulations) on each time point moving backwards from the maximum difference to identify the earliest timepoint that showed a statistically significant difference (*p* < .05) with consecutive significant timepoints leading up to the maximum difference (Wessel & Aron, 2015; Wessel & Huber, 2019).^1^

Bayesian Wilcoxon signed-rank tests indicated moderate evidence that the peak burst latency following unexpected haptic events in the CMO task occurred earlier than after the Stop signal on successful Stop trials in the SST (*BF_10_* = 6.26). However, despite numerically earlier peak burst latencies, there was moderate evidence for no difference between unexpected auditory events and the Stop signal (*BF_01_* = 4.84) and anecdotal evidence for no difference between unexpected visual events and the Stop signal (*BF_01_* = 1.77). In Dataset 2, strong evidence indicated that the peak burst latencies following both unexpected auditory and unexpected visual events preceded the peak burst latency for the Stop signal (Unexpected Auditory: *BF_10_* = 1.48×10^4^, Unexpected Visual: 19.81). Strong evidence indicated that the onset of the FC P3 (isolated from the Stop signal IC) following each type of unexpected event occurred earlier than that for the Stop signal (Dataset 1, Unexpected Auditory: *BF_10_* = 204.19; Unexpected Visual: *BF_10_* = 273.86, Unexpected Haptic: *BF_10_* = 36.56; Dataset 2, Unexpected Auditory: *BF_10_* = 133.14; Unexpected Visual: *BF_10_* = 6423.26). In the Discussion, we speculate on the possibility that inhibitory processes may be accelerated following unexpected events in settings that do explicitly require inhibition.

When comparing the peak burst latencies to the FC P3 onsets, strong evidence indicated that the peak burst latency following unexpected haptic events preceded the corresponding onset of the FC P3 (*BF_10_* = 673.10), but such evidence was lacking for the other types of unexpected events (Unexpected auditory: *BF_01_* = 1.93, Unexpected Visual: *BF_01_* = 5.12). In the SST, the peak burst latency on successful Stop trials also preceded the corresponding P3 onset (*BF_10_* = 7.15). In Dataset 2, the peak burst latency following unexpected auditory events preceded the corresponding P3 (*BF_10_* = 1.58×10^5^), although moderate evidence was found for no difference for unexpected visual events (*BF_01_* = 5.10). Once again, the peak burst latency on successful Stop trials preceded the corresponding P3 onset (*BF_10_* = 3.06). Overall, these findings provide some support for the notion that FC β-bursts precede the FC P3. This was most apparent within the SST. It bares mentioning that these tests may have been conservative for potential timing differences given that peak burst latency was compared to a measure of the onset of the FC P3. Nevertheless, the combined evidence across tasks and datasets suggest that FC β-bursts reflect an earlier occurring signature than the FC P3.

## DISCUSSION

We found clear and consistent support for our prediction that FC β-bursts would be increased following unexpected events. In Dataset 1, which included unexpected events in auditory, visual, and haptic modalities, increased β-bursting at frontal electrode sites was evident in all three modalities. In Dataset 2 (from Wessel & Huber, 2019), which included unexpected events in the auditory and visual modalities, increased FC β-bursting was evident following unexpected auditory events, and, numerically, following unexpected visual events as well. To our knowledge, this represents the first investigation of β-bursts following unexpected events.

Jones and colleagues have recently suggested a role for β-bursts in the detection of sensory events. In particular, Shin et al. (2017) showed that β-bursts over somatosensory cortices originating 300 to 100 ms before tactile stimulus delivery impaired perception whereas Law et al. (2022) showed that β-bursts contemporaneous with stimulus delivery aided perception. While we did not explicitly explore the consequences of β-bursts on perception, our results suggest that FC β-bursts may also play a role in stimulus detection and perhaps in registering their surprise value. The finding of early increased FC β-bursting following unexpected events compared to standard events already suggests that these bursts may play a role in the detection of infrequent events. However, our finding that these bursts also scale with Shannon surprise on just the subset of trials with unexpected events joins and expands upon this finding to suggest that this infrequency-representation happens at very fine-grained scale (capturing trial-to-trial deviations in expected probabilities).

Across both datasets, FC β-bursting was linked to a specific type of surprise: Shannon surprise with separate terms for each modality. Although correlated, Shannon surprise and Bayesian surprise differ mainly in that an unexpected event can have a higher surprise value than its predecessor with a sufficiently long stretch of intermediate expected events according to the Shannon model. Our experimental design did not explicitly attempt to dissociate the two types of surprise (which has been previously attempted e.g, in O’Reilly et al., 2013). Given that Bayesian surprise and Shannon surprise were correlated in the datasets, however, it is even more interesting that FC β-bursts specifically correlated with Shannon surprise in both cases, and to a consistently lesser extent with Bayesian surprise. It is unlikely that this dissociation is explainable in terms of differing degrees of sensitivity to detect either relationship as both types of surprise were predictive of RT and the whole-brain EEG data corresponding to the FC P3 response at the single-trial level (for analyses relating FC P3 to surprise, see Appendix). Prior work has indicated that Shannon surprise and Bayesian surprise relate to at least somewhat distinct neural processes. For instance, using fMRI, O’Reilly et al. (2013) identified activity within posterior parietal cortex as representing Shannon surprise whereas activity in anterior cingulate cortex and pre-supplementary motor area uniquely represented Bayesian surprise (i.e., updating, as they refer to it). Scalp-surface EEG studies suggest that whereas FC P3 better characterizes Bayesian surprise, Shannon surprise is better characterized by centro-parietal P3 (Mars et al., 2008; Kolossa et al., 2015; Seer et al., 2016; Wessel & Huber, 2019). Interestingly, the present results suggest that FC β-bursts might represent a unique frontal cortex neural correlate of Shannon surprise and one that emerges sooner than the P3 components. To our knowledge, this represents the first demonstration of a signature of Shannon surprise over frontal cortex. Presumably, this early detection of infrequent events or surprise could serve to pave the way for rapid adjustments to behavior and cognition (O’Reilly et al., 2013; Wessel & Aron, 2017; Diesburg & Wessel, 2021).

The time course of this increase in FC β-bursts and its relation to surprise was further interesting. Although the sliding window approach we employed to analyze β-bursts still introduces temporal uncertainty as to the precise timing with which increased β-bursting begins, we could at least be confident that FC β-bursts were elevated soon after the presentation of the unexpected event and well before 200 ms. This is important in establishing FC β-bursts as a potentially early signature because, notably, the putative onset of the FC P3 process within the context of action stopping occurs subsequently, around 200 ms (Wessel & Aron, 2015). At the group-level, increases in FC β-bursting appeared to occur well before the onset of the FC P3 in both the CMO task and the SST (see Fig. 8). By adopting an individual subject-level approach to comparing peak burst latency and FC P3 onset, we were further able to show that peak β-bursting following unexpected haptic events in Dataset 1 and unexpected auditory events in Dataset 2 preceded the onset of the corresponding P3. While we did not find evidence for differences with the other unexpected events, it is worth noting that comparing the peak of the β-burst process to the onset of the P3 may be quite conservative. Moreover, peak β-bursting following the Stop signal preceded the onset of the corresponding P3 in both datasets. These findings offer some initial insight into the potential processes that the FC β-burst signature might relate to within a couple of our theoretical frameworks. Wessel and Aron’s (2017) adaptive re-orienting theory of attention and Diesburg and Wessel’s pause-then-cancel model (2021, see also: Schmidt & Berke, 2017) commonly propose multi-stage models whereby an early stage is characterized by global inhibition of the motor system brought about by the detection of unexpected or otherwise salient, behaviorally relevant events which is then followed by a later stage in which more strategic adjustments and updates to ongoing motor and cognitive programs occur. Based on the timing of various functional neurophysiological signatures and their relation to behavior, Diesburg and Wessel (2021) have proposed certain functional signatures to be associated with either the purported early or late inhibitory stages. For instance, studies involving single-pulse TMS stimulation of M1 regions responsible for muscles irrelevant to the task at hand both within action stopping contexts (i.e., the SST or its variants) and within following unexpected events have estimated global inhibition to begin around 150 ms, thus providing the main functional signature of the pause phase. Other signatures such as brief decreases in EMG around this time (Jana et al., 2020; Raud & Huster, 2017; Raud et al., 2020; Tatz et al., 2021) or the still earlier, transitory decreases in isometric force contractions (Novembre et al., 2018) also likely relate to this stage. By contrast, the main purported neurophysiological correlate of the cancel phase is the FC P3 signature, which has been tied to successful response inhibition in the SST and RT slowing following unexpected events. However, a direct neural correlate of the pause phase with precise timing such as EEG or magnetoencephalography is lacking. Moreover, based on previously observed timings of increased frontal β-bursting following the stop signal (Jana et al., 2020; Wessel, 2020), it remained unclear which processing stage this inhibitory signature related to and whether the increases in frontal β-bursting index reactive inhibitory control or might instead relate to proactive inhibitory control. In the primarily reactive context of the CMO task, elevated FC β-bursts emerged soon after event onset and would overall seem better aligned with the timing of the pause phase than the cancel phase. In the SST involving the same individuals, elevated β-bursting emerged somewhat later (∼178 ms) but still prior to 200 ms and the emergence of the stop-signal P3. This timing is rather intermediate to the ∼120 ms timing found by Jana et al. (2020) and the ∼200 ms timing found by Wessel (2020), but nevertheless shifts the evidence in favor of a timing more consistent with the pause phase.

We also found that when isolating the FC P3 from the Stop signal IC, the onsets of the FC P3 occurred earlier following each type of unexpected event in the CMO task than following the Stop signal on successful Stop trials in the SST. We also found peak β-burst latencies to occur earlier following unexpected haptic events than on successful Stop trials in Dataset 1. We did not find earlier burst latencies following unexpected auditory or visual events in Dataset 1; however, we did in Dataset 2. Thus, an intriguing possibility is whether surprise accelerates the inhibitory process. Some initial evidence for this comes from Iacullo et al. (2020) who examined cortical-spinal excitability while participants responded to imperative stimuli that were quickly followed by standard tones, infrequent/expected tones, or novel/unexpected sounds. Critically, they found global motor suppression to occur earlier following novel/unexpected sounds compared to infrequent/expected tones. In contrast to surprising events, the SST contains not just the reactive inhibitory control brought about by the infrequent stop-signal itself, but also proactive inhibitory control, or strategic adjustments in anticipation of the stop-signal (Elchlepp et al., 2016; Wessel, 2018b). To better capture proactive control, researchers have created variants of the SST that include certain go trials (in contrast to the default, maybe stop trials). However, fMRI studies that have done so have implicated BOLD responses in the same regions (Chikazoe et al., 2009; Jahfari et al., 2010), and EEG studies have likewise implicated some of the same event-related potentials, including FC P3, in both types of control (Elchlepp et al., 2016; Raud & Huster, 2017). Soh et al. (2021) found increased sensorimotor β-bursts on maybe stop compared to certain go trials, a difference that was present even before trial onset. Interestingly, Muralidharan et al.’s (2022) recent work modeling frontal β-bursts in the SST suggested that these bursts increased the decisional threshold necessary for the Go response. Taken together, studies on the SST suggest that some of the same processes subserve proactive and reactive inhibitory control. If so, it would seem plausible that these processes might take place more rapidly when proactive control is minimal, as in the case of surprising events. From the standpoint of rapidly allocating resources and adjusting behavior, this would indeed be an adaptive feature as, at least in most real-world settings, the optimal course of action (or inaction) cannot be known in advance of the surprising event but instead must be urgently gleaned after its occurrence. For example, after detecting something suddenly crawling across their skin, one might decide to hold still. Unexpected events may therefore provide a framework that is better aligned with how inhibition transpires in our day to day (Wessel, 2018a). Overall, the current results are consistent with the notion that inhibitory processes may be accelerated with surprise. We note, though, that while all unexpected events prompted earlier FC P3 onset latencies compared to those evoked by the Stop signal, the results concerning FC β-bursts were more mixed (i.e., with three of the five unexpected event conditions showing earlier latencies than on successful Stop trials). Most basically, we herein demonstrate that frontal β-bursts occur in reactive contexts.

In the present task designs, FC β-bursts were not directly related to the observed motor slowing. One possibility is that β-bursts reflect an early processing stage that only indirectly relates to RT. Within the proposed pause phase (Diesburg & Wessel, 2021), information must pass through cortex and multiple subcortical structures before arriving at primary motor cortex. By contrast, the proposed cancel phase is argued to more directly relate to actual behavioral adjustments. Subsequently, the correspondence of FC β-bursts with RT may have been too subtle to detect a relationship (even as RT is a highly variable measure itself), even though this lack of relationship could still be consistent with pause phase roles of salient event detection or otherwise triggering inhibition. In addition, the FC P3, which is a prominent signature of the cancel phase, exhibited correspondences with RT following each type of unexpected event (see *Fig.* A2). Another reason for not finding a connection between FC β-bursts and RT could owe to the nature and timing of our CMO task. In our task, unexpected events occurred as a part of a cue that occurred 500 ms before the imperative stimulus. Thus, any early inhibition prompted by the unexpected events might have been overcame by the time the stimulus appeared. Rather, the timing of our task may have been more optimal for capturing the impact of the cancel phase on inhibition, and indeed we found relationships between RT and FC P3. In hindsight, the timing of increased FC β-bursting might have also occurred too soon. For instance, Little et al. (2019) showed that movement was only disrupted by sensorimotor β-bursts that occurred just before the response. Here, responses occurred even sometime after target onset (as participants complete a choice RT task). Given that these issues could have compounded in the present task design, it is interesting to consider whether a relationship between β-bursts and behavior might be established in a task in which unexpected events occur during movement which could be assessed for instance by presenting unexpected events just after the first increase in electromyographic activity. Regardless, if global inhibition serves to promote rapid action adjustments in the first place, it would seem plausible that this process might be more active when actions are already underway.

In sum, we have shown that unexpected events in three modalities (auditory, haptic, and visual) prompt increases in FC β-burst rate soon after their onset and before 200 ms. These β-bursts further appear to track the surprise (or trial to trial fluctuations in stimulus infrequency) of these events separately for each modality and as indexed by Shannon surprise. Although we were unable to establish a relationship between FC β-bursts and RT, the overall results suggest that FC β-bursts may play an early role in inhibitory processes such as salient event detection and/or the triggering of the global inhibition characteristic of the pause phase.

## Data Availability Statement

All data and analysis scripts will be deposited publicly on the OSF upon acceptance of the manuscript.

## Acknowledgements

We thank Saara Engineer, Nathan Chalkley, Nathan Chen, and Kylie Dolan for their help with data collection. We thank Benjamin Rangel for assistance with the set up and monitoring of the startle response in pilot participants before and after experimentation. We thank two anonymous reviewers for suggesting additional supporting analyses.

## Author contributions

Joshua R. Tatz: Conceptualization; Formal analysis; Investigation; Visualization; Writing – Original Draft, Writing – Review & editing. Alec Mather: Conceptualization; Investigation; Formal analysis. Jan R. Wessel: Conceptualization; Formal analysis; Funding acquisition; Investigation; Project administration; Supervision; Writing – Review & editing.

## APPENDIX

As a subsidiary goal of this study, we sought to identify whether the EEG data in Dataset 1 replicated the whole-brain event-related EEG findings reported by Wessel and Huber (2019), especially since their data were re-analyzed as Dataset 2. We also aimed to extend their findings to unexpected events originating from the haptic modality. In general, the results closely followed those of Wessel and Huber (2019) further underscoring the similarity of Datasets 1 and 2, despite the inclusion of the third haptic modality in Dataset 1.

## EEG analyses following Wessel and Huber (2019)

We considered four results from Wessel and Huber (2019) to be of primary importance in replicating their work:

1. single-trial EEG data should favor a model with separate Bayesian surprise terms in all three modalities (auditory, visual, and haptic)
2. mean RT should be predicted by FC P3 in each modality
3. the FC P3 following unexpected events should be recoverable from a FC IC isolated from the stop-signal task when used as a functional localizer (i.e., the stop-related FC P3)
4. the separate Bayesian surprise model should be implicated with this IC alone

Given that we examined β-bursts for both Shannon and Bayesian surprise, for the sake of thoroughness, we also newly examined the abovementioned analyses in Dataset 2 based on Shannon surprise. The ensuing description of the methods relevant to this subsection is adapted from Wessel and Huber (2019):

First, the surprise values originating from both types of surprise (Shannon, Bayesian surprise) and modality considerations (separate, combined) were used to model the whole-brain event-related single-trial EEG response on trials with unexpected events. We based this analysis on procedures reported by Fischer and Ullsperger (2013). To compare surprise models, at the single-trial level, we used robust regression [in MATLAB, robustfit()] to fit the standardized (*z*-scored) EEG signal with the standardized (*z*-scored) surprise values in each participant in each electrode in each trial in 48 ms time windows spanning from 50 to 500 ms. These data were baseline corrected with the data from −100 to 0 ms and averaged across trials (48 trials, barring those excluded because of artifacts). For each model specification, this resulted in a 64 (channel)* 10 (time point) = 640 matrix of beta weights. At the group level, the mean beta weights corresponding to the models were tested against 0 (using one-sample *t*-tests) and separate and combined models were tested against each other (using paired-samples *t*-tests) within each type of surprise. Multiple comparisons were corrected for three sets of 640 *t*-tests via a false discovery rate correction (FDR-corrected, *p* < .01) procedure (FDR, Benjamini et al., 2006).

Second, we examined Bayesian Pearson correlations between the mean amplitudes of the P3 responses following each type of unexpected event (auditory, visual, and haptic) and the corresponding mean RT. The subject-average amplitudes were determined by finding the largest positive-going amplitude deflection in the trial-average (at electrodes FCz and Cz) during the time-window ranging from 150-500 ms after the unexpected event (or in the case of the visual P3, we modified this search range to 250-600 ms based on the characteristics of the response reported in Wessel and Huber).

Third, the SST was employed as a functional localizer to demonstrate that the surprise-related FC P3 response shares a common neural generator with the inhibition related FC P3 response in the SST. To this end, the two datasets from the CMO and SST task were merged, and the preprocessing steps and the ICA detailed above were repeated anew on the merged datasets. The stop-related P3 IC was selected automatically using a two-step spatiotemporal selection procedure (Wessel & Ullsperger, 2011). First, components were identified whose back-projected channel-space topography showed a maximum positive difference in stop compared to go trials in a FC ROI (consisting of electrodes FC1, FCz, FC2, C1, Cz, and C2 from among 9 ROIs across the scalp). From these components, the component that showed the highest correlation to the original channel-space ERP in the same FC ROI and time window was selected for each participant as their stop-related IC. In Dataset 1, we validated the IC selected stop-signal P3 using a two-fold procedure that addresses whether the stop-signal P3 occurs earlier on successful stops and whether its onset is positively correlated with SSRT. In short, we replicated prior work using this validation procedure (Kok et al., 2004; Wessel & Aron, 2015; Wessel & Huber, 2019). We do not report these results here (but refer the interested reader to the data and code on OSF).

Fourth, we repeated the surprise model analyses detailed in (1) using only each subject’s selected Stop-related IC for the unexpected event in the CMO task. As control analyses, we subjected two other versions of the merged dataset to the same analyses. These included reconstructions of the channel-space based on a back-projection of all original ICs *sans* the Stop-related IC and a back-projection of the IC that accounted for the most residual variance after the Stop-related IC was removed. As in (1), these tests were FDR-corrected for multiple comparisons (which again included 640 tests per set).

## Replication of Wessel and Huber (2019) and extension to haptic domain

We now report on the four results that we considered to be of primary importance (as detailed in the corresponding portion of the Method section) to replicate Wessel and Huber (2019) and extend their findings to the haptic domain:

First, using Dataset 1, we aimed to replicate Wessel and Huber’s (2019) finding involving Bayesian surprise that a model with separate surprise terms for each modality would better explain the single-trial EEG response than the combined data and that the spatiotemporal dynamics of any significant time periods would relate to those of the P3 response. These tests were FDR-corrected for a family-wise alpha-level of .01. As can be seen in Fig. A1, the separate surprise term model showed a significant, positive prediction of the EEG data beginning with the 326-374 ms bin and extending through the remaining time bins (i.e., to 476-524 ms) to around the onset of the imperative stimulus. For the combined surprise term model, the 326-374 ms bin also showed a significant positive prediction of the EEG data which also extended through the end of the trial. However, the separate surprise term model fit the data better than the combined surprise term model beginning with the 326-374 ms bin and onwards. Compared to Wessel and Huber, the bin in which we first observed the separate surprise term model was slightly later (326-374 ms compared to 276-324 ms). This might be attributable to the larger sample size in Dataset 2 or the inclusion of unexpected events in the haptic modality in Dataset 1. Nevertheless, this time range coincided with the FC P3 response within each modality. In addition, the mean standardized beta-weights at model-fitting electrodes identified by Wessel and Huber (FC2, C1, Cz, and C2) were all reliably above zero when fitting the Bayesian surprise model with separate modality terms. We therefore replicated Wessel and Huber with the current, trimodal dataset in showing that a model with separate surprise terms (involving Bayesian surprise) predicts the single-trial EEG data.

**Fig. A1.**
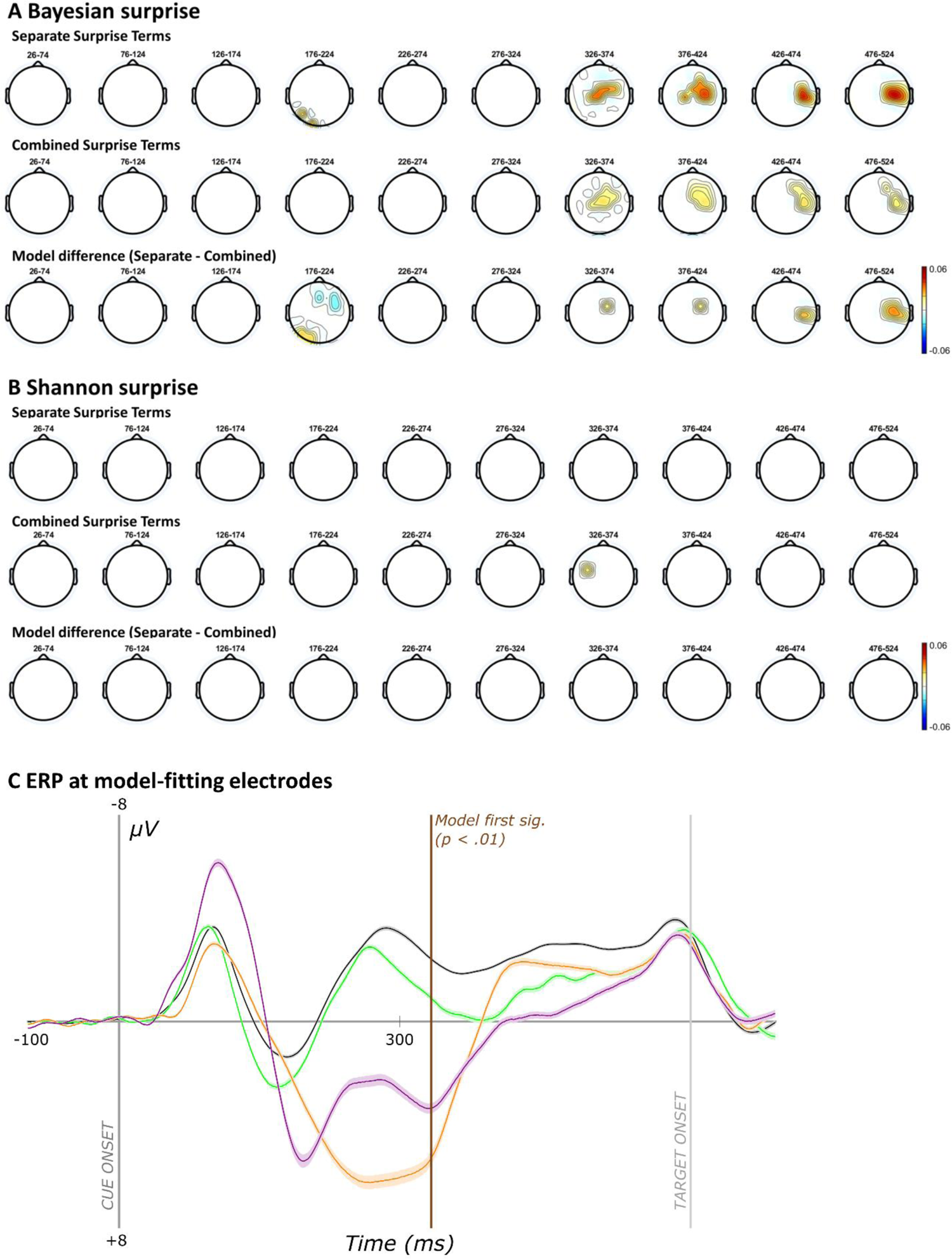
Scalp-topographies from modeling whole-brain single-trial EEG data with surprise and event-related potential at model-fitting electrodes in Dataset 1. **a** Topographies represent the mean standardized individual beta weights for Bayesian surprise and separate modality terms (top row), Bayesian surprise and combined modality terms (mid row), and the difference in fits between these two models (bot row). Heatmaps reflect beta weights. **b** The same analyses as panel a but with Shannon surprise terms. c Cue-locked event-related potentials following each event type at model-fitting electrodes (FC2, C1, Cz, and C2) that were significant for the winning Bayesian surprise model and in Dataset 2. The spatiotemporal dynamics of the model-fitting electrodes correspond to the FC P3.

Given that we examined β-bursts with respect to Shannon surprise (in addition to Bayesian surprise), we repeated the above analyses using Shannon surprise and with both datasets (since Wessel and Huber had not examined this for Dataset 2). For Dataset 1 (see Fig. A1b), Shannon surprise fitted the whole-brain EEG data at some electrode sites (both positively and negatively), but rather inconsistently and not at FC sites. For Dataset 2, however, Shannon surprise significantly fit the data in the 326-374 ms time bin and at FC sites. Significant model fits were not found with common surprise terms and no differences between model fits were found, either.

Second, we correlated the mean amplitude of the FC P3 response in each modality with mean RT (see Fig. A2). Medium-sized positive correlations were found between RT and P3 amplitude following unexpected events in each modality, including the new haptic modality (Unexpected visual: *r* = .39, *BF_10_* = 8.18; Unexpected audio: *r* = .37, *BF_10_* = 5.35; Unexpected haptic: *r* = .32, BF_10_ = 2.45). This is consistent with Wessel and Huber’s findings in the auditory and visual modalities and further links this functional signature to inhibition (as larger responses brought about slower RTs).

**Fig. A2.**
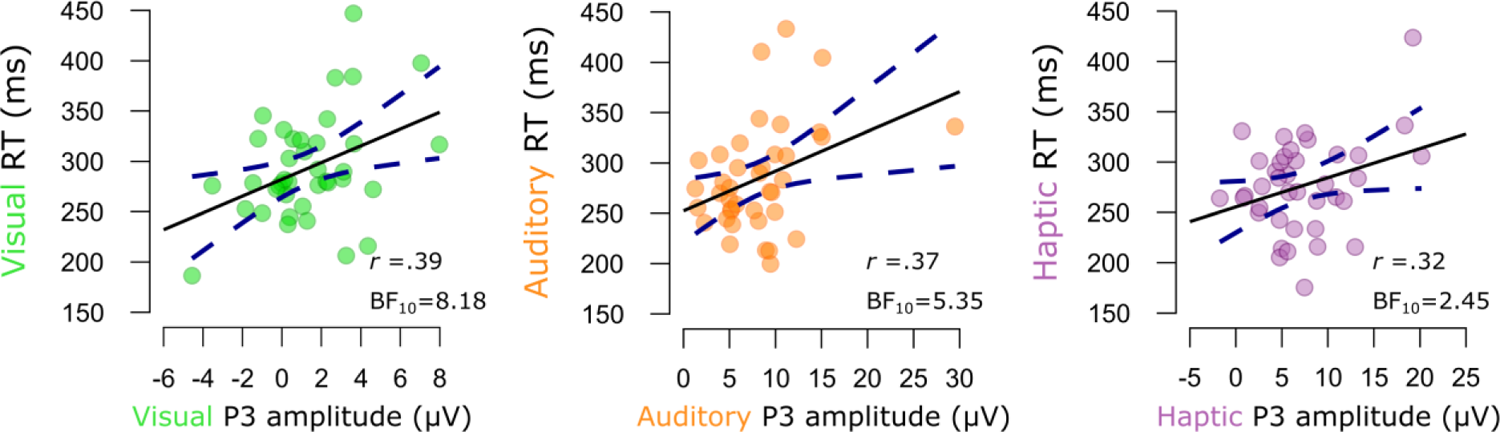
Pearson correlations between mean FC P3 amplitude and mean RTs following each type of unexpected event in Dataset 1. Dots indicate the means from individual subjects. Solid black lines indicate the slope of the linear model. Dotted dark blue lines indicate the ±95 % confidence intervals.

Third, having merged the datasets from the visual SST and CMO tasks and having isolated a frontocentral stop-related IC from each subject, we aimed to recover the FC P3 response using this one independent component. Since ICA serves to unmix a signal into its constituent (orthogonal) sources, identifying the FC P3 in the CMO task by using the SST as a functional localizer would suggest that a common neural generator is responsible for the FC P3 in both contexts. Since the SST was completed only in the visual modality, identification of FC P3 in the other auditory and haptic modalities would further highlight this signature as a domain-general inhibitory signature—granted, one that still retains information about the sensory modalities prompting inhibition (otherwise, the surprise model fitting results would have favored the common surprise term model). The difference waves are shown in Fig. 8 in the main text. We were indeed able to produce the FC P3 in the CMO task when using the SST as a functional localizer task.

**Fig. A3.**
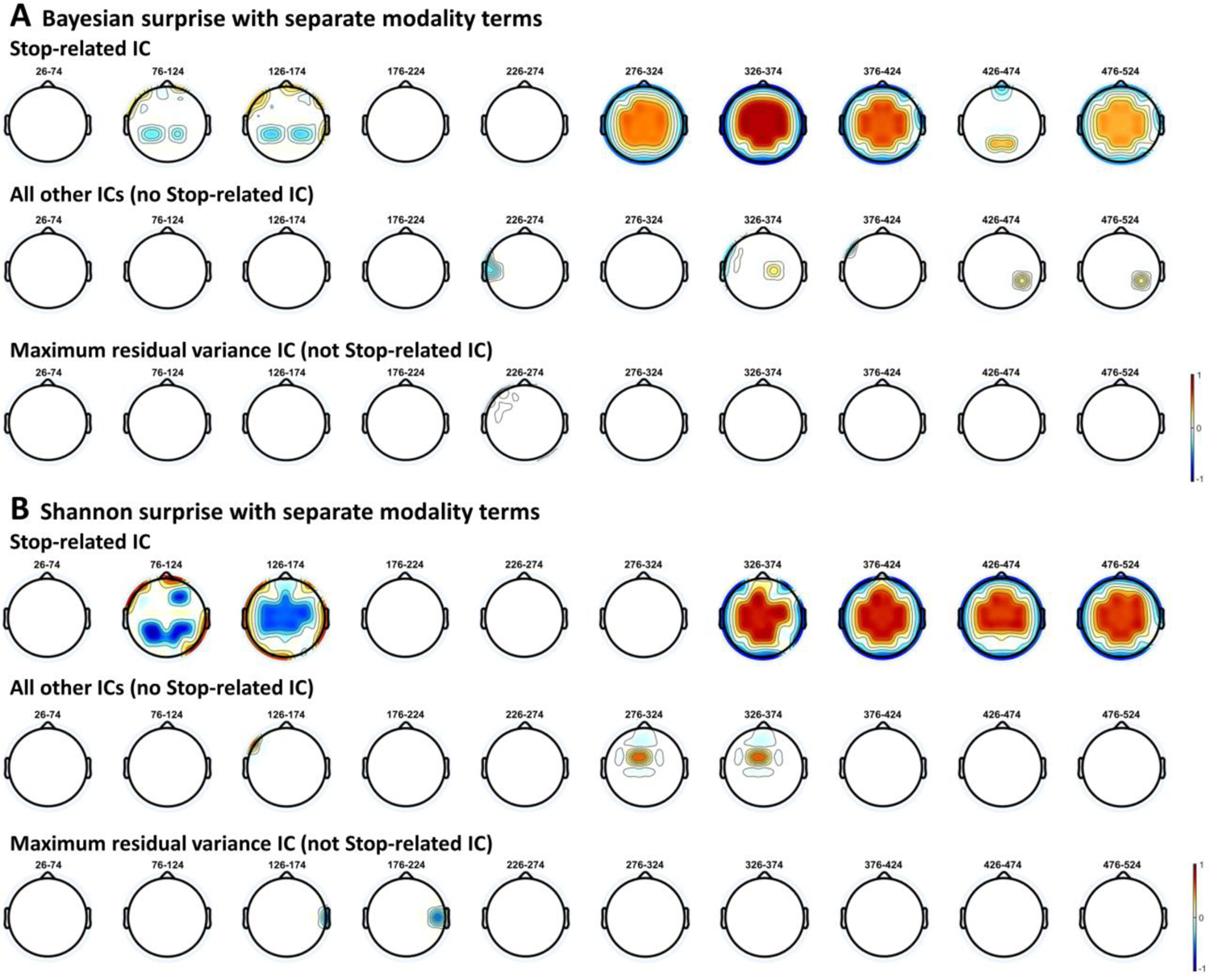
Scalp-topographies of the ICs modeled with Bayesian surprise and Shannon surprise in the CMO task of Dataset 1 (n=39). **a** Beta-weights from the single-trial model-fitting of IC-EEG data with Bayesian surprise under the winning separate modality term model. Colors indicate significant model fits (warm colors = positive, cool colors = negative; white = non-significant, p = .01, FDR-corrected). **b** The same analyses but with Shannon surprise.

**Fig. A4.**
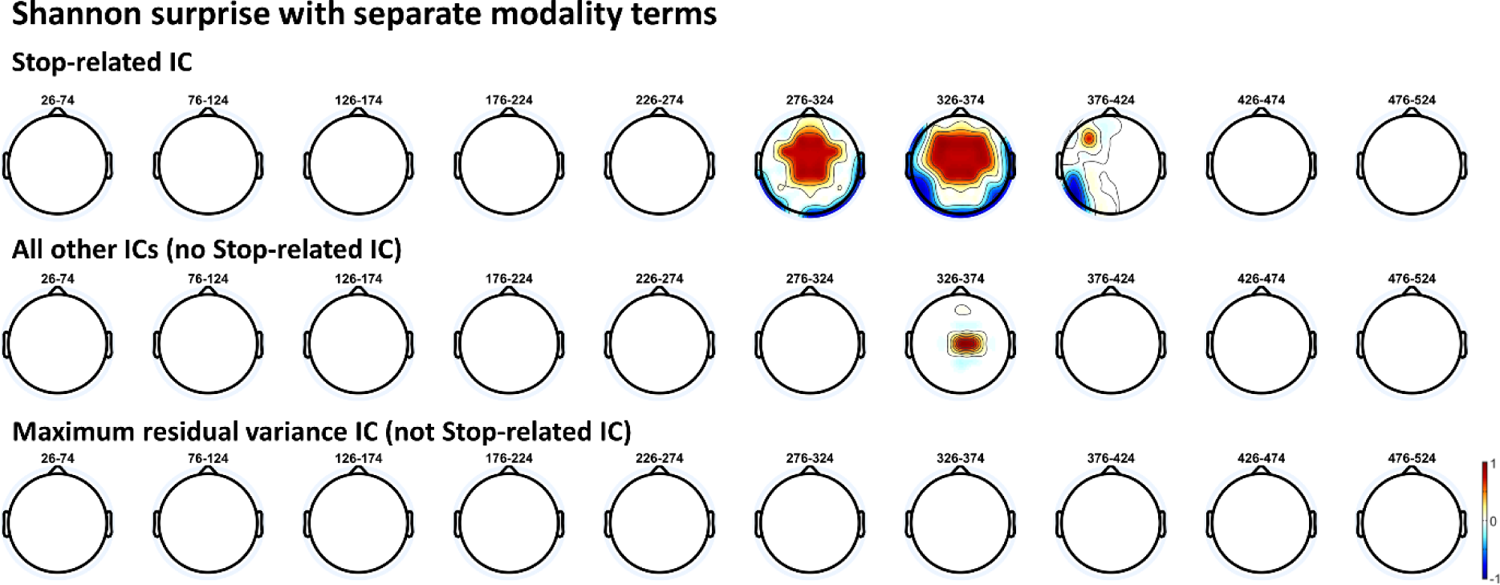
Scalp-topographies of the ICs modeled with Shannon surprise in the CMO task of Dataset 2 (n=55). The same as Fig. A3b but with Dataset 2.

As our final replication of Wessel and Huber with Dataset 1, we repeated the abovementioned whole-brain, single-trial EEG response modeling with surprise, this time using only the Stop-related IC. For control analyses, we also completed these analyses using all other ICs and the IC that explained the most variance upon removal of the stop-related IC. As can be seen in Fig. A3a, Bayesian surprise clearly fit the stop-related IC in Dataset 1. Moreover, this occurred at FC sites and timepoints consistent with the FC P3, suggesting that the neural generator involved in action inhibition in the SST also tracked surprise within the context of the CMO task. Interestingly, with the stop-related IC, Shannon surprise also fit the data at similar electrode sites and timepoints (although Shannon surprise became significant at FC sites in the 326-374 ms as opposed to the 276-324 ms which was significant with Bayesian surprise). This contrasts the model-fitting analysis involving the full EEG data which did not show this relationship to Shannon surprise (see Fig. A1). Dataset 2 also replicated this finding of positive relationships between Shannon surprise and stop-related IC activity at FC sites. Here, significant model fits were found in the 276-324 and 326-374 ms bins. Most basically, the results from the IC-based approach replicate Dataset 2 in suggesting that the FC P3 process involved in action stopping also tracks surprise within the context of a CMO task. We further show here that using the IC-based approach, Shannon surprise also predicts activity at FC sites around the time of the FC P3 in Dataset 2 (see Fig. A4). This could simply be because our experimental designs did not attempt to decorrelate Bayesian surprise and Shannon surprise. Future work might attempt to do so by intentionally introducing long stretches of intervening standard events between the same type of unexpected events to see if FC P3 is better captured by Bayesian surprise and whether another IC relating to the centro-parietal P3 better captures Shannon surprise (e.g., Mars et al., 2008; Seer et al., 2016).

## Footnotes

1 We note that using a similar procedure to identifying onsets on the β-burst data returned earlier differences than the peak in only a handful of participants, we therefore only considered peak burst latency in the ensuing analyses.

